# Two temperature-responsive RNAs act in concert: The small RNA CyaR and the mRNA *ompX*

**DOI:** 10.1101/2024.10.08.617194

**Authors:** David A. Guanzon, Stephan Pienkoß, Vivian B. Brandenburg, Jennifer Röder, Alisa Dietze, Andrea Wimbert, Christian Twittenhoff, Franz Narberhaus

## Abstract

Bacterial pathogens, such as in *Yersinia pseudotuberculosis*, encounter temperature fluctuations during host infection and upon return to the environment. These temperature shifts impact RNA structures globally. While previous transcriptome-wide studies have focused on RNA thermometers in the 5’-untranslated region of virulence-related mRNAs, our investigation revealed temperature-driven structural rearrangements in the small RNA (sRNA) CyaR. At 25°C, CyaR primarily adopts a conformation that occludes its seed region, but transitions to a liberated state at 37°C. By RNA-sequencing and in-line probing experiments, we identified the Shine-Dalgarno sequence of *ompX* as a direct target of CyaR. Interestingly, the *ompX* transcript itself exhibits RNA thermometer-like properties, facilitating CyaR base pairing at elevated temperatures. This interaction impedes ribosome binding to *ompX* and accelerates degradation of the *ompX* transcript. Furthermore, we observed induced proteolytic turnover of the OmpX protein at higher temperatures. Collectively, our study uncovered multi-layered posttranscriptional mechanisms governing *ompX* expression, resulting in lower OmpX levels at 37°C compared to 25°C. We propose that CyaR remains mostly passive at 25°C but becomes primed to repress *ompX* at 37°C, thereby expediting outer membrane remodelling in anticipation of pathogenesis.

## INTRODUCTION

Bacteria are masters of adapting to an ever-changing environment by modulating gene expression at every level of gene regulation, spanning from transcription to translation and beyond. Regulatory RNAs are key mediators of posttranscriptional control [1]. Positioned strategically within the 5’-untranslated region (5’-UTR) of the mRNA are RNA thermometers (RNATs) [2, 3] and riboswitches [4]. These *cis-*regulatory RNA elements affect gene expression by remodeling the secondary structure upon ligand binding (riboswitches) or in response to temperature fluctuations (RNATs). These structural rearrangements, in-turn, regulate processes such as transcription termination or translational initiation [5].

In addition to these localized elements, small regulatory RNAs (sRNAs) can act as both *cis*- and *trans*-regulatory elements to fine-tune cellular responses, in particular in stress pathways [6]. Varied in size (ranging from 50 to more than 200 nucleotides) and complexity of secondary structure [7] sRNAs feature distinctive domains such as internal U-rich sequences upstream of factor-independent terminators (Class I sRNAs), ARN motifs (Class II sRNAs) [8], and 3’ poly-U tails that facilitate interaction with the RNA chaperone Hfq [9, 10]. This protein plays a pivotal role in maintaining intracellular sRNA stability in many bacteria [8, 10] and in the recognition of target transcript through the formation of ternary sRNA-Hfq-mRNA complexes [11]. In contrast to *cis-*regulatory sRNAs, which possess regions fully complementary to their target mRNAs, *trans*-regulatory sRNAs find and affect their target transcripts through limited base pairing through a so-called seed region [12]. In principle, the accessibility of the seed region is critical for the establishment of sRNA-mRNA interactions [13, 14]. Base-paired regions between sRNAs and mRNAs vary, ranging from large, predominantly complementary pairings of ∼30 bp seen in MicF-*ompF* [15] and Spot42-*galK* [16], to imperfect interactions of approximately 13 bp with bulges as observed in SdsR-*ompD* [17], to very short (≥7-bp) yet multiple binding domains as in RyhB with various *omp* transcripts [14], to as few as 6-bp in SgrS-*ptsG* [18]. This modular design enables numerous sRNAs to orchestrate extensive regulatory networks [19].

In a limited number of examples, *cis*-acting riboswitches or RNATs have been shown to work in concert with *trans*-acting sRNAs. For instance, in *Listeria monocytogenes*, virulence gene expression is regulated by an RNAT in the 5’-UTR of *prfA*, which encodes the virulence master regulator. Two *trans*-acting sRNAs, termed SreA and SreB, originating from S-adenosylmethionine riboswitches, bind to this 5’-UTR of *prfA*, establishing a connection between bacterial virulence and nutrient availability [20]. In *Escherichia coli*, the sRNAs OmrA and OmrB interact with the adenosylcobalamin riboswitch present in the *btuB* mRNA and thereby repress synthesis of the vitamin B12 importer [21]. In enterohemorrhagic *E. coli*, the sRNA CyaR interacts with the RNAT in the *chuA* mRNA, which encodes an outer membrane haem receptor [22].

In our investigation, we have uncovered a pair of temperature-responsive RNAs that coordinate gene expression at multiple levels. Specifically, we found a regulatory interplay between the sRNA CyaR and the mRNA *ompX* in *Yersinia pseudotuberculosis*. This food-borne pathogen employs a diverse array of riboregulatory mechanisms to sense the temperature transition from the external environment to the host milieu, thereby redirecting bacterial metabolism towards pathogenicity [23]. First and foremost, *lcrF* coding for the master regulator of virulence genes is under RNAT control [24]. However, many other genes associated with virulence and metabolic processes are also regulated by RNATs as determined by transcriptome-wide RNA structure profiling both *in vitro* [25] and *in vivo* [26]. Thermo responsive structural RNA elements were found upstream of genes coding for a secreted toxin [27], the various components of the type 3 secretion system (T3SS) [28, 29], an outer membrane protein potentially implicated in virulence [30], and critical enzymes in the oxidative stress response [31].

Interestingly, our global RNA structure probing studies have revealed temperature-responsive structural motifs not only in mRNAs but also in sRNAs, among them in CyaR [25, 26]. Originally identified as RyeE in *E. coli* [32], this sRNA was later renamed to CyaR (<u>Cy</u>clic AMP-<u>a</u>ctivated <u>R</u>NA) upon discovery of its regulation by Crp [33]. As a highly conserved sRNA among enterobacteria, the regulatory networks of CyaR have been well-studied in *E. coli* [34–36] and *Salmonella* [13]. In *Yersinia*, although CyaR remains under the control of Crp, its expression pattern extends beyond the availability of carbon sources and late growth phases to changes in temperature [37]. Our RNA structuromics datasets [25, 26] suggest that CyaR undergoes dynamic structural rearrangements, particularly within its seed region in response to temperature elevation. Consequently, it is plausible to speculate that the melting of the sRNA structure during *Yersinia’s* transition from the external environment to the host conditions, might increase its regulatory capacity by liberating the seed region and facilitating target recognition.

In this study, we demonstrate that the sRNA CyaR in *Y. pseudotuberculosis* undergoes temperature-dependent structural remodeling. We further establish that this structural adaptation enhances its regulatory efficacy, particularly on the *ompX* mRNA, which harbours its own temperature-responsive RNA structure. As a consequence of translational control, mRNA stability and protein stability mechanisms, we observe a cumulative effect on OmpX levels, which are lower at 37°C compared to 25°C.

## MATERIALS AND METHODS

### Bacterial strains and growth conditions

Strains used in this study are listed in table S1. Bacteria were grown aerobically in LB medium (1% tryptone, 1% NaCl, and 0.5% yeast extract) at 25°C and 37°C, unless otherwise indicated. When necessary, supplements and antibiotics were used at the following concentrations: ampicillin 150 µg/ml, chloramphenicol 20 µg/ml, kanamycin 50 µg/ml, *Yersinia* specific antibiotics (cefsulodin 15 µg/ml, triclosan 4 µg/ml, novobiocin 2.5 µg/ml), spectinomycin 300 µg/ml, rifampicin 250 µg/ml, sucrose 10% w/v, L-arabinose 0.2% v/v.

### Plasmids and strain construction

Plasmids and oligonucleotides are listed in tables S.1 and S.2, respectively. For standard cloning methods or gene assemblies, NEBuilder HiFi DNA Assembly Kit (New England Bio Labs, Frankfurt, Gemrany) was utilized following the manufacturer’s protocols in the construction of the plasmids.

The plasmid pBAD2-*bgaB* served as the vector backbone for BgaB fusion constructs. The 5’-UTR up to 30 nucleotides of the coding sequence of *ompX* (from -43 to +30 from AUG) was amplified with (OmpX_BgaB_Fw/OmpX_BgaB_Rv) primer pair and digested with NheI and EcoRI, then ligated into vectors linearized with the same restriction enzymes generating plasmid (pBO6899).

For GFP reporter gene assays, the gene assembly consisted of the plasmid pXG-10 [38] linearized via PCR using the oligonucleotides (pXG_UTRs_Fw/pXG_Gibson_Rv) and a 152 bp PCR amplified fragment using oligos (pXG_OmpX_gib_Fw/pXG_OmpX_Rv) corresponding to the 5’-UTR up to 45 nucleotides of the coding sequence to generate (pBO7535). Control plasmid pXG0 was generated by linearizing pBO7535 with oligonucleotides (pXG_Ev_Gibson_Fw/pXG_Ev_Rv) to recreate the empty vector (pXG0), and pXG1 was generated by linearizing pBO7535 with oligonucleotides (pXG1_GFPEv_Fw/pXG1_GFPEv_Rv) to remove the insert and leave the GFP with its own RBS and start codon, and re-circularized using KLD enzyme mix (New England Bio Labs, Frankfurt, Germany).

Run-off transcription plasmids containing a primer-encoded upstream T7 promoter were constructed for *in vitro* RNA synthesis for RNA structure probing and primer extension inhibition. CyaR was amplified with the oligos (CyaR_T7_Fw/CyaR_EcoRV_Rv), and *ompX* was amplified with (OmpX_rnf_Fw/OmpX_rnf_Rv). The fragments were then blunt-cloned into a SmaI digested pUC18 plasmid to generate (pBO5019 & pBO7904) respectively.

For site-directed mutagenesis, oligonucleotides encoding the desired mutations were used to amplify the plasmids carrying the WT constructs of *ompX* RNAT and re-circularized using KLD enzyme mix (New England Bio Labs, Frankfurt, Germany) to generate the following variants: *ompX* R1 (UU_23-24_CC) (OmpX_WT_SD_Fw/OmpX_Rep1_Rv) generating the following plasmid (pBO8202), *ompX* R2 (UU_23-24_CC; G_28_C) (OmpX_R2_G28C_Fw/OmpX_Rep1_Rv) generating the following plasmid (pBO8203). CyaR stable (UUU_82-83,87_GGC) and CyaR open (UUC_84-86_AAA) were synthesized (Thermo Scientific, Waltham, USA) to obtain the following plasmids (pBO7428 & pBO7430).

To generate a polar deletion mutant of CyaR, a mutant allele where CyaR was replaced with a kanamycin resistance cassette (*neoR*) flanked by a 510 bp homologous region upstream and a 407 bp homologous region downstream was cloned into the suicide plasmid pYD32 (pGP704) and was a kind gift from the Dersch lab [39]. To chromosomally integrate the mutant allele, *Y. pseudotuberculosis* YPIII (recipient strain) was conjugated with an *E. coli* S17-1 *λ-pir* (donor strain) carrying the plasmid. Single crossover mutants were selected with LB plates containing *Yersinia* specific antibiotics and chloramphenicol. Putative deletion mutants that underwent double crossover events were counterselected for *sacB* with sucrose. The deletion mutants were further validated with PCR and DNA sequencing using primers (CyaR_internal_Fw/CyaR_internal_Rv & CyaR_external_Fw/CyaR_external_Rv).

To complement the deletion mutant, 89bp fragment encoding for CyaR was amplified using the primers (CyaR_NcoI_Fw/CyaR_PstI_Rv) and digested with NcoI and PstI, then ligated into the arabinose inducible vector pGM930 linearized with the same restriction enzymes to generate (pBO5009). To construct chromosomal *ompX^3xFLAG^,* a 1346 bp fragment was amplified corresponding to 782 bp upstream of, and 564 bp downstream of the transcription start site using the primer pair (OmpX_SacI_Fw/OmpX_XbaI_Rv). This fragment was then digested with SacI and XbaI, then ligated into pDM4 suicide vector digested with the same enzymes to generate (pBO6885). To integrate the tagged version of *ompX, Y. pseudotuberculosis* YPIII (recipient strain) was conjugated with an *E. coli* S17-1 *λ-pir* (donor strain) carrying pBO6885. Single crossover mutants, where pDM4 was chromosomally integrated, were selected with LB plates containing *Yersinia* specific antibiotics and chloramphenicol.

### RNA isolation and Northern blot analysis

Total bacterial RNA was isolated from cell pellets using hot acid phenol method and performed as described previously [28, 40]. Transcripts were then detected via Northern blot performed as described in earlier works [41]. *Y. pseudotuberculosis* genomic DNA was used to amplify DNA templates used to generate DIG-labeled RNA probes. Gene specific forward oligos were paired with gene specific reverse oligos with an upstream T7 promoter, primer pairs for CyaR: (CyaR_probe_Fw/CyaR_probe_Rv) and *ompX*: (OmpX_probe_Fw/OmpX_probe_Rv) to generate complementary RNA strands during *in vitro* transcription using DIG-labeled uridine residues. An *in vitro* prepared DIG-labeled *gfp* RNA probe was generated as described previously [28].

### RNA half-life

To observe the regulatory efficiency of CyaR as it transitions different temperatures, cells were initially grown at 25°C to an OD_600_ 0.5. The cells were then split to a batch that remained at 25°C and the other half was transferred into prewarmed flasks at 37°C. Rifampicin was immediately added to abolish transcription. Samples were taken at the indicated timepoints and snap frozen in liquid nitrogen, and relative transcript amounts were analyzed by Northern blot. Densitometry measurements were done with ImageLab v6.1 (BioRad) software.

### RNA sequencing

RNA levels were analyzed at an OD of 0.5 and 1.5 at 25 and 37 °C from strains *Y. pseudotuberuclosis* YPIII and ΔCyaR grown in LB medium. Purified RNA was isolated and treated with the TURBO DNA-free Kit™ (Thermo Scientific, Waltham, USA) according to the manufacturer’s instructions to avoid DNA contamination. Library preparation and RNA-sequencing (Illumina NovaSeq 6000 platform) was performed by Novogene Co., LTD. Data analysis was performed as described in [42] with the difference that an adjusted p-value of 0.01 and a twofold up-or downregulation relative to the wild type were defined for identification of differentially expressed genes. Putative sRNA-mRNA interactions were predicted by the IntaRNA algorithm by setting the minimum number of base pairs to five nucleotides [43]

### Reporter gene assay

For cells carrying *bgaB* fusions of *ompX*, cells were grown in LB with ampicillin at 25°C up to an OD_600_ of 0.5, then transcription was induced with L-arabinose. The cells were split, where half of the culture was transferred into prewarmed flasks at 37°C and the other half remained at 25°C and incubated a further 30 mins. Samples were then taken for β-galactosidase assay and Western blot analysis. As positive control, *bgaB* fused with the short 5’-UTR of *yopN* plus 30 nt of the CDS was used [28]. Specific activities of the enzyme were then calculated based on previous work [44] and presented as mean ± SD of three biological replicates.

For cells carrying *gfp* fusions, cells were grown in LB with chloramphenicol at the indicated temperatures and harvested at the indicated OD_600_. Total bacterial RNA was then isolated and *gfp* transcripts were then detected by Northern blot analysis. GFP fluorescence measurements were done by growing the cells in LB with chloramphenicol in a black, clear-bottom 96-well plates (Nunc, Thermo Scientific) (Starting OD_600_ 0.01). Cell density was monitored by measuring OD_600_ (600 nm) and GFP fluorescence intensities was measured by (Wavelength Excitation: 480 nm / Emission: 520 nm) in a 96-well plate reader (Infinite M Plex, Tecan) at the indicated temperatures. To calculate the absolute fluorescence, cellular autofluorescence measurements from cells carrying an empty vector was subtracted from measurements from cells carrying *gfp* fusion plasmids. Relative fluorescence was then calculated by dividing absolute fluorescence values by the OD_600_. For fold changes, relative fluorescence values in the wild type strain carrying the *ompX::gfp* fusion was set to 1 and presented as mean ± SD of three independent cultures.

### Western Blot analysis

Whole-cell protein samples were prepared to visualize the temperature-dependent regulation of OmpX. Briefly harvested cells were resuspended in 1x SDS sample buffer (2% (w/v) SDS, 12.5 mM EDTA, 1% (w/v) β-mercaptoethanol, 10% (v/v) glycerol, 0.02 % (w/v) bromphenol blue, 50 mM Tris, pH 6.8) and protein extracts were adjusted to 0.01 OD_600_/µl. Whole-cell protein extracts were then denatured at 95°C for 10 min.

To visualize OmpX::3xFLAG expression, total proteins were separated in 12.5 % SDS polyacrylamide gels. They were then blotted onto nitrocellulose membranes and stained with Ponceau S to assure equal loading of proteins. To detect OmpX::3-FLAG, α-FLAG primary antibody (1:4000, Sigma-Aldrich) was used followed by a goat α-Mouse IgG (H+L)-HRP conjugate (1:4000, BioRad, DE). Signals were detected with Immobilon Forte Western HRP substrate (Merck, DE) and (ChemiDoc^TM^ MP, BioRad, DE).

For quantitative analysis of expression, whole-cell proteins were prepared as described above but resolved in 12% TGX stain-free gels (BioRad, USA) then activated under UV-light for 45 sec prior to transfer onto a nitrocellulose membrane. Similarly, an α-FLAG primary antibody (1:4000, Sigma-Aldrich) was used followed by goat α-mouse IgG Starbright Blue 700 (1:2500, BioRad,DE) protected from light. After six 5-minute washes with 1x TBST, the membranes were then dried for 15 min, prior to detection (ChemiDoc^TM^ MP, BioRad, DE). Densitometry measurements were then done with ImageLab v6.1 (BioRad) software.

### Protein half-life

To observe if OmpX regulation extends up to the protein level, cells were grown at 25°C or 37°C. When indicated, the cells were split to a batch that remained at 25°C and the other half was transferred into prewarmed flasks at 37°C or vice versa. Spectinomycin was immediately added to stop translation. Samples were taken at the indicated timepoints and snap frozen in liquid nitrogen. The relative protein levels were analyzed by quantitative western blots.

### Structure probing of RNAs

To probe the secondary structures of RNA at different temperatures, EcoRV linearized plasmids were used as DNA templates for *in vitro* transcription using the T7 polymerase (Thermo Scientific, Waltham, USA) as described previously [28].

For enzymatic structure probing, the plasmids (pBO5019, pBO7428, and pBO7430) were used to synthesize RNA for the entire transcript of CyaR WT along with the stable and open variants, respectively. The transcripts were then gel-purified in 6% polyacrylamide gels, dephosphorylated, and the RNA was 5’-labeled with [^32^P] as described previously [45]

Single-hit kinetics for the radio-labeled RNA (30,000 cpm) was achieved by adding ribonuclease T1 (0.0016 U) (Thermo Scientific, Waltham, USA) or T2 (0.0025 U) in 5xTN buffer (500 mM NaCl, 100 mM Tris acetate, pH 7) and incubating at 25, 37, and 42°C for 5 minutes. Reactions were stopped with formamide stop solution.

For in-line structure probing, 5’-[^32^P] labeled *ompX* transcript from (pBO7904) was prepared as described above. Single-hit kinetics for the radio-labeled RNA (30,000 cpm) was achieved by incubating equal volumes of labeled RNA suspended in RNase free water and 2x In-line probing buffer [5] (100 mM Tris-HCl pH 8.3, 200 mM KCl, 40 mM MgCl_2_), when indicated CyaR transcripts were added, adjusting water volumes to compensate for the additional transcript. The mixture was then incubated in a thermocycler (CFX, BioRad, USA), with heated lids to prevent evaporation, samples at 25°C were incubated for 40 hrs and samples at 37°C were incubated for 10 hrs. Reactions were stopped with formamide stop solution.

For the alkaline ladder, 60,000 cpm of labeled RNA was used and incubated with 1 µL of 10x ladder buffer (1 M Na_2_CO_3_, 1 M NaHCO_3_, pH 9) for 2 minutes at 90°C. For the T1 ladder, 30,000 cpm of labeled RNA was used and incubated at 90°C with 1 µL of sequencing buffer (provided with the RNase T1) to fully denature the RNA, then incubated with the T1 nuclease for an additional 5 min at 37°C. The samples were separated on an 8% polyacrylamide gel. Phosphorscreens were then detected by GE Amersham^TM^ Typhoon^TM^ (GE Healthcare, USA). Densitometry measurements were done with ImageLab v6.1 (BioRad) software.

### Primer extension inhibition assay

*In vitro* transcribed *ompX* were synthesized from the same plasmid (pBO7904) mentioned in structure probing. 5’-[^32^P]-labeled reverse primer (OmpX_rnf_Rv). 30S ribosomal RNA and tRNA^fMet^ (Sigma-Aldrich, St. Louis, USA) were used for the assay as described previously (Hartz D, Extension inhibition analysis of translation, 1988). Briefly, an annealing mix was prepared consisting of 0.16 pmol radiolabeled primer and 0.08 pmol RNA mixed with 1x VD-Mg^2+^ (60 mM NH_4_Cl, 6 mM β-mercaptoethanol, 10 mM Tris/HCl, pH 7.4) and denatured for 3 mins at 80°C and annealed at -20°C for 30 mins. To bind the 30S ribosomal subunit and CyaR, 16 pmol of tRNA^fMet^, 0.16 pmol radiolabeled primer, 6 pmol 30S ribosomal subunit in Watanabe buffer (60 mM HEPES/KOH, 10.5 mM Mg(CH_3_COO)_2_, 690 mM NH4COO, 12 mM β-mercaptoethanol, 10 mM spermidine, 0.25 mM spermine), and 1.5 pmol of CyaR were mixed, then incubated at 10 mins at 25, 37, and 42°C. For the extension reaction, an M-MLV mix (1x VD+Mg^2+^, (10 mM Mg(CH_3_COO)_2_, 6 µg BSA, 4 mM dNTPs, 800 U M-MLV reverse transcriptase (Thermo Scientific, Waltham, USA)) was added to initiate the cDNA synthesis for 10 mins at 37°C. The reactions were stopped with formamide stop solution and separated on an 8% polyacrylamide gel. For orientation, a sequencing ladder was generated with the Thermo Sequenase sequencing kit (Thermo Scientific, Waltham, USA) following the manufacturer’s instructions.

## RESULTS

### The seed region of CyaR but not the 5’-end is conserved

A genome-wide search in *E. coli* led to the identification of highly conserved small RNAs, including CyaR (originally known as RyeE) [32]. Sequence alignments of CyaR across *E. coli, Salmonella* and various *Yersinia* species reveal a remarkable conservation in the seed region (Fig. 1A), while the 5’ region exhibits considerable variability in accordance with the phylogenetic relationships of the bacteria (Fig. S1). Secondary structure predictions at both 25 and 37°C using RNAfold indicate the presence of a short thermo-responsive element within the initial ten nucleotides of *Yersinia* CyaR, a feature that is absent in the corresponding RNAs from *Salmonella*, *Escherichia* and *Shigella* (Fig. S2). Furthermore, CyaR from these organisms displays a different overall structure in comparison to their *Yersinia* homologs. *In silico* analysis predicts the seed region to adopt a short unresponsive hairpin structure across all examined organisms, including *E. coli* and *Salmonella* CyaR, which have been previously characterized [13, 33].

**Figure 1.**
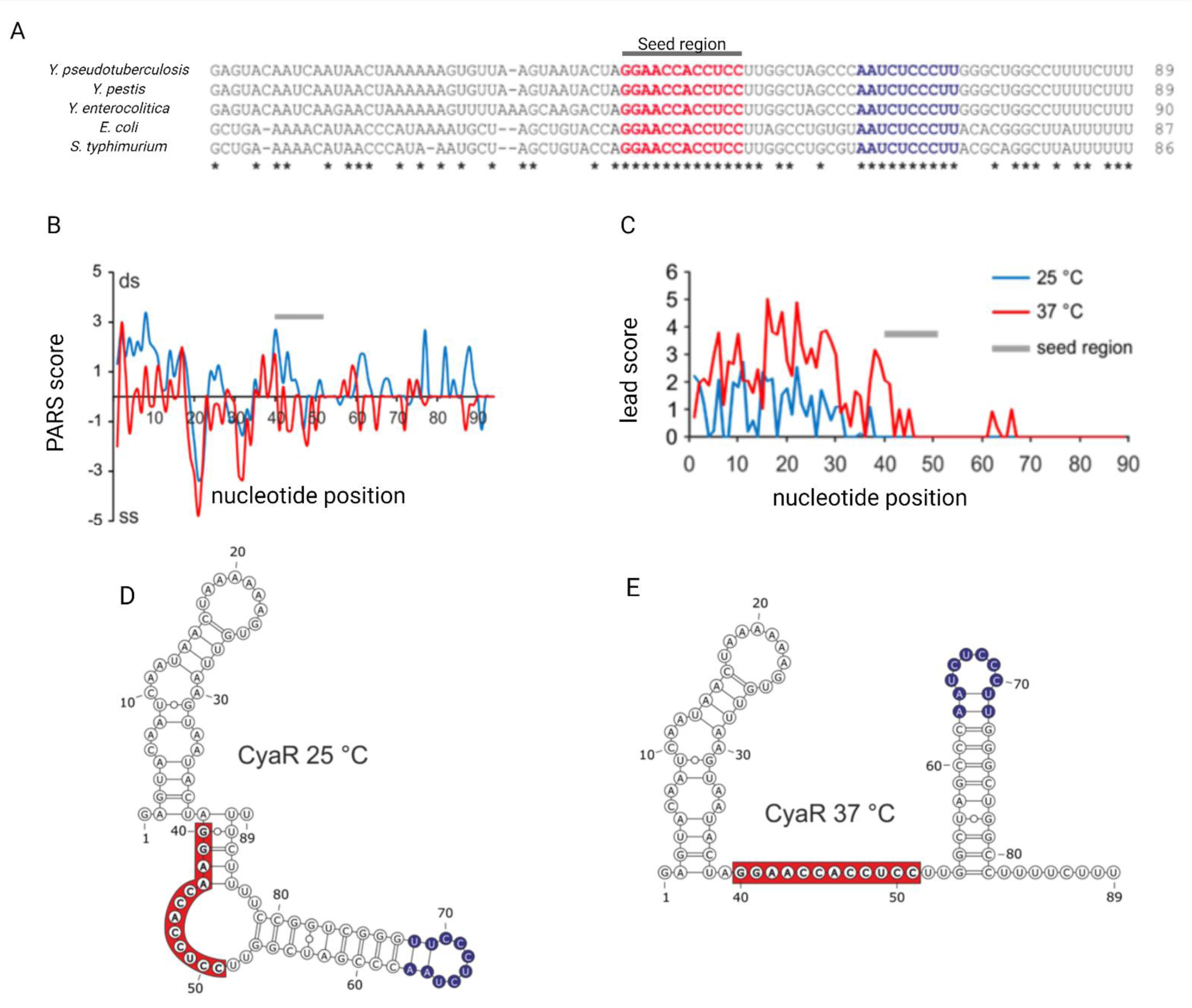
*Yersinia*-specific sequences in *Yersinia* CyaR contribute to the unique thermo-responsive structure. (A) RNA sequence alignment of selected closely related species. (B) PARS profiles of *Y. pseudotuberculosis* CyaR at 25°C and 37°C [25]; ds, double stranded; ss, single-stranded. (C) Lead-seq profile of *Y. pseudotuberculosis* CyaR at 25°C and 37°C [26]. The lead score reflects the single-strandedness of each nucleotide. (D, E) PARS-derived RNA secondary structures at 25°C and 37°C. Highlighted in red is the highly conserved seed region and blue is the conserved sequence in the terminator loop.

### Temperature-dependent structural remodelling of CyaR

In contrast to the computational RNA structure predictions, our experimental findings derived from transcriptome-wide RNA structuromics studies in *Y. pseudotuberculosis* indicate (i) a distinctive CyaR structure, and (ii) a temperature-dependent structural rearrangement, both *in vitro* [25] and *in vivo* [26]. The Parallel Analysis of RNA Structures (PARS) profiles delineate double-stranded and single-stranded nucleotides, which were mapped *in vitro* using RNase V2 and RNase S1, respectively (Fig. 1B). Our results reveal that nucleotides 39 to 43 in the presumed seed region are base-paired at 25°C and transition into a single-stranded conformation at 37°C. Intriguingly, a similar change was observed at the 3’ end of CyaR, as indicated by a transition from a positive to a negative PARS score at 37°C. The RNA structures inferred from these global probing results suggest that the seed region becomes occluded by the 3’ poly-U tail (84-UUCUU-88) at 25°C, with this base pairing unwinding at 37°C (Fig. 1 D-E). Furthermore, the results from the lead-seq experiment [26], which identified lead-sensitive, single-stranded nucleotides *in vivo*, largely support this model. Specifically, the seed region remained protected from lead acetate-induced cleavage at 25°C, while becoming accessible to cleavage at 37°C (Fig. 1C).

To complement the findings derived from transcriptome-wide RNA structure probing approaches, we elucidated the structure of *in vitro*-synthesized CyaR at 25, 37 and 42°C (Fig. 2). We employed RNase T1, which cleaves single-stranded RNA at the 3’ end of guanines and RNase T2, known for introducing cuts preferentially at the 3’ end of adenines but also at other positions. The outcome of the T2 experiment confirmed the presence of loop regions spanning nucleotides 19-23 and 65-68, which were susceptible to cleavage by the RNase at all three temperatures (Fig. 2A and B). These regions are devoid of guanosines and thus remained unaffected by the RNase T1. Nucleotide A30 and its adjacent nucleotides were susceptible to RNase cleavage across all temperatures indicating an accessible conformation. Of particular interest, the nucleotides forming base pairs in the predicted seed region of CyaR (40-GGAA-43) were barely cleaved at 25°C but became increasingly susceptible to T1 and/or T2 at elevated temperatures. These observations are consistent with the secondary structures calculated from PARS analysis, as depicted in Fig. 1D and E.

**Figure 2.**
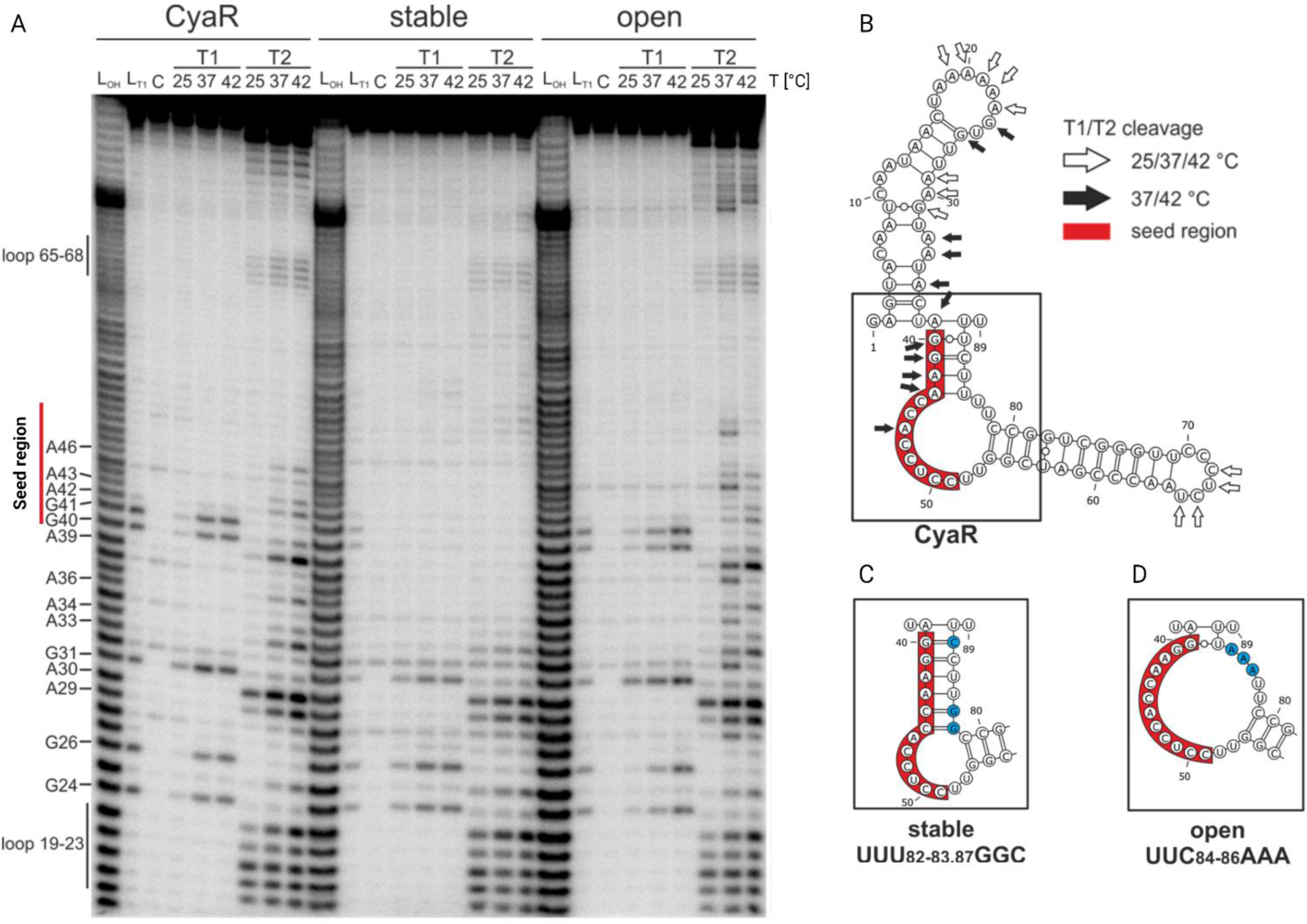
The putative CyaR seed region is occluded at 25°C *in vitro.* (A) Enzymatic structure probing of the thermo-responsive structure of CyaR WT, and stable and open variants at 25, 37, and 42°C. 5’ ^32^P-labeled RNA was treated with single-strand specific RNases T1 (0.0016 U) and T2 (0.0025 U) at the indicated temperatures. From left to right: L_OH_: Alkaline ladder; L_T1_: T1-treated RNA at 37°C as reference for G residues; C: water control. (B) PARS-derived secondary structure of CyaR WT at 25°C. White arrows indicate cleavage at all temperatures tested. Black arrows indicate cleavage only at 37 and 42°C. (C, D) RNAfold-predicted secondary structures of stabilized and destabilized CyaR variants. Nucleotide substitutions are depicted in blue. The enzymatic structure probing gel is a representative of two independent experiments.

Rationally designed nucleotide exchanges were introduced in the 3’ end of CyaR (UUU_82,83,87_GGC; Fig. 2C) to strengthen the interaction with the seed region. As a result, the seed region become entirely shielded from cleavage by both T1 and T2 RNases (Fig. 2A). Mutations intended to weaken the interaction between the seed region and the 3’end of CyaR (UUC_84-86_AAA, Fig. 2D) resulted in a WT-like cleavage pattern (Fig. 2A), suggesting the persistence of residual RNA-RNA interactions.

Based on these cumulative structure probing results (Fig. 1 and 2), we formulated the hypothesis that the seed region of CyaR becomes more accessible at elevated temperatures, which, in turn, facilitates more efficient regulation of its mRNA targets.

### CyaR is a CRP- and Hfq-dependent small RNA in *Y. pseudotuberculosis*

Before moving on to the identification of CyaR target transcripts, we aimed to learn more about CyaR expression in *Y. pseudotuberculosis*. Previous Northern blot experiments had revealed consistently high levels of CyaR during the stationary growth phase at both 25 and 37°C [37]. Our own Northern blots corroborated these findings (Fig. 3A to C). In addition, we found a growth-phase dependency of CyaR levels at 25°C, with its levels increasing towards the stationary phase. At 37°C, CyaR levels remained elevated throughout both low and high optical densities.

**Figure 3.**
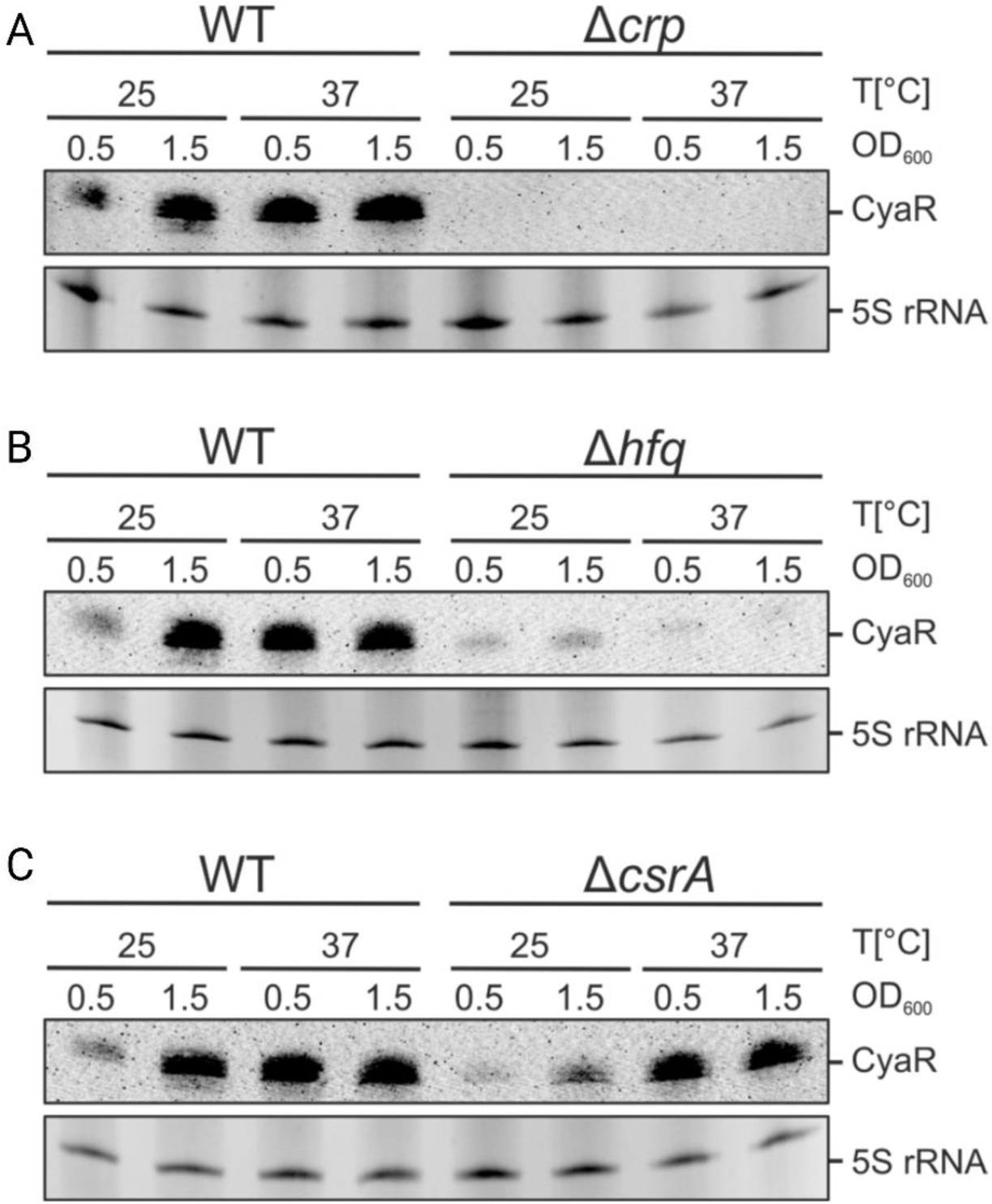
*Y. pseudotuberculosis* CyaR is CRP-and Hfq-dependent. (A to C) CyaR expression in the wildtype (WT) and various deletion strains under different temperature conditions and growth phases. The Northern blots shown are representatives of three independent experiments.

Previous electrophoretic mobility shift assays had demonstrated the binding of the CyaR promoter by the global regulator CRP [37]. Our findings further validate the CRP-dependence of CyaR in *Y. pseudotuberculosis*, as evidenced by the absence of detectable CyaR transcripts in the *crp* mutant across all four tested conditions (Fig. 3A). Furthermore, we found a critical dependence of CyaR levels on the presence of Hfq (Fig. 3B) but not on CsrA (Fig. 3C)

### RNA-sequencing revealed 63 differentially regulated genes in ΔCyaR

To identify potential mRNA targets of CyaR in *Y. pseudotuberculosis*, we constructed a strain lacking CyaR by substituting the genomic sRNA locus with a kanamycin resistance cassette. We verified the correct identity of this mutant by PCR (Fig. S3A), followed by DNA sequencing. Due to its overlapping genomic location, we simultaneously deleted the putative sRNA transRNA_24, annotated on the complementary strand of CyaR. However, the expression of transRNA_24 remained undetectable under any of the conditions relevant to this study (Fig. S3B), strongly indicating that all observed effects in the CyaR mutant were solely attributable to the absence of CyaR.

We conducted a comparative RNA-seq analysis of the WT strain and the ΔCyaR mutant cultured in LB medium under four different conditions: early and late exponential growth phase, and temperatures of 25 and 37°C. Our analysis revealed a total of 63 genes exhibiting either up- or downregulation of at least twofold in the ΔCyaR mutant under at least one of the tested conditions (Fig. 4A). Among these, the most prominent target was *ompX*, a well-established negatively regulated CyaR target in *E. coli* and *Salmonella* [13, 33, 34]. In *Y. pseudotuberculosis*, the *ompX* mRNA was significantly upregulated in the CyaR mutant across three conditions, coinciding with the high abundance of the sRNA in the WT strain under these conditions (compare Fig. 3A-C). Although not meeting the cutoff of two-fold induction, *ompX* expression followed the same trend at 25°C in early exponential growth phase, showing a modest 1.77-fold induction in the CyaR mutant (Fig. 4B). The observed differential expression of *ompX* at 25°C contradicts our initial hypothesis, which proposed that CyaR might primarily interact with target mRNAs under elevated temperatures due to conformational changes. This finding suggests that the regulatory mechanism governing *ompX* expression by CyaR is more complex than anticipated.

**Figure 4.**
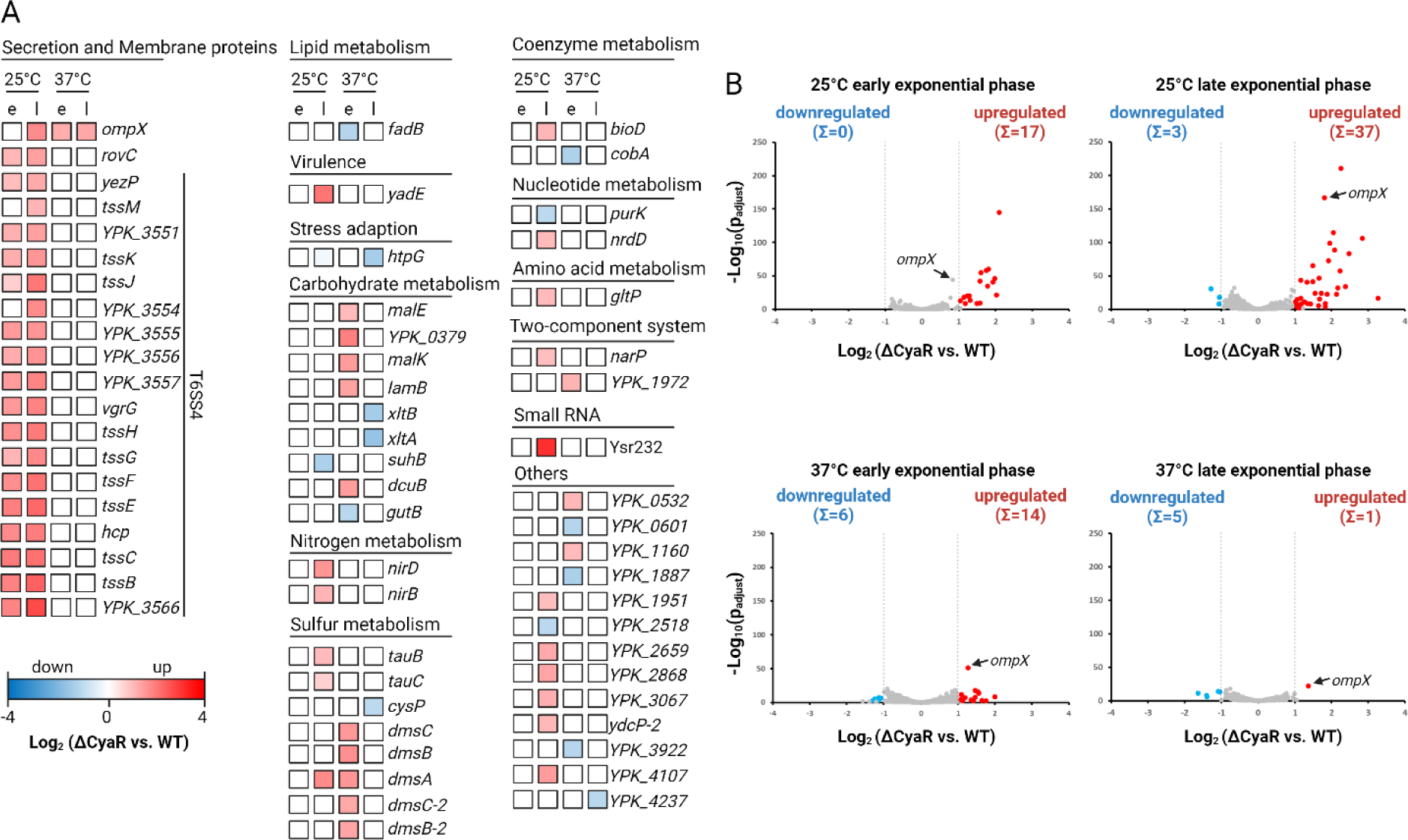
RNA sequencing reveals putative targets of CyaR in *Y. pseudotuberculosis.* (A) Genes were categorized based on their annotated function, and further grouped according to their regulation under specific conditions, i.e. at 25 or 37°C and/or at the early (e) or late (l) exponential phase. Levels of regulation are represented in a heatmap with color-coded boxes. Blue boxes denote downregulation, whereas red boxes indicate upregulation. The gradient of Log_2_ fold changes is shown in bottom left corner. -(B) Volcano plots visualize genes differentially expressed in the ΔCyaR strain versus the WT at the indicated conditions. Genes were considered differentially expressed in the transcriptome analysis (n= 3 biological replicates) if cut-off values set at: adjusted p-value ≤ 0.001, Log_2_ Fold Change ≥ 1-fold-change(s) up or down were met.

Other genes differentially expressed in the CyaR mutant are involved in secretion and metabolic pathways of carbohydrates and inorganic compounds (Fig. 4A). Overall, a total of 49 genes were upregulated, whereas 14 genes were downregulated indicating a predominantly negative regulatory role of CyaR on potential mRNA targets (Fig. 4B).

Remarkably, the entire T6SS4 gene cluster coding for a type VI secretion system, along with *yezP* encoding a known T6SS4 effector, and *rovC* coding for the cognate transcriptional regulator, were all upregulated in the ΔCyaR mutant at 25 °C. However, the normalized read counts and the differences observed between the WT and ΔCyaR strain were relatively low, typically in the low hundreds (for example, a difference of approximately 280 reads for *YPK_3567* (*rovC*) between the strains). This expression level falls below the detection limit of Northern blot analysis for validation experiments. Furthermore, it is important to consider that the expression of the *Y. pseudotuberculosis* T6SS4 system is typically weak or even silent under normal laboratory conditions, as it is tightly regulated [46]. Hence, we are uncertain whether the observed differences between the WT and CyaR are physiologically meaningful.

### Alternative methods confirm CyaR-mediated regulation of *ompX* expression

For further analysis, we selected *ompX* as a prime candidate due to its pronounced differential regulation under all tested conditions (Fig. 4), coupled with sufficiently high transcript levels to permit detection via Northern blot analysis. First, we evaluated the levels of chromosomally encoded *ompX* mRNA in *Y. pseudotuberculosis* WT and CyaR mutant, both carrying empty vectors, along with the CyaR deletion strain complemented with the CyaR plasmid at both low and high cell densities (Fig. 5A). At 25°C, the absence of CyaR resulted in an increase in *ompX* transcripts, suggestive of negative regulation by the sRNA in the WT. While the restoration of normal *ompX* levels by a 10-min pulse induction of CyaR was less efficient in the early growth phase, it was highly effective in the later growth phase. At 37°C, *ompX* transcripts were barely detectable in all three strains. However, the overall trend, with slightly elevated *ompX* levels in the ΔCyaR strain, mirrored that observed at 25°C. The generally low level of *ompX* transcripts at 37°C suggest the involvement of additional transcriptional and posttranscriptional regulatory mechanism, for example by other sRNAs, like MicA [33, 47].

**Figure 5.**
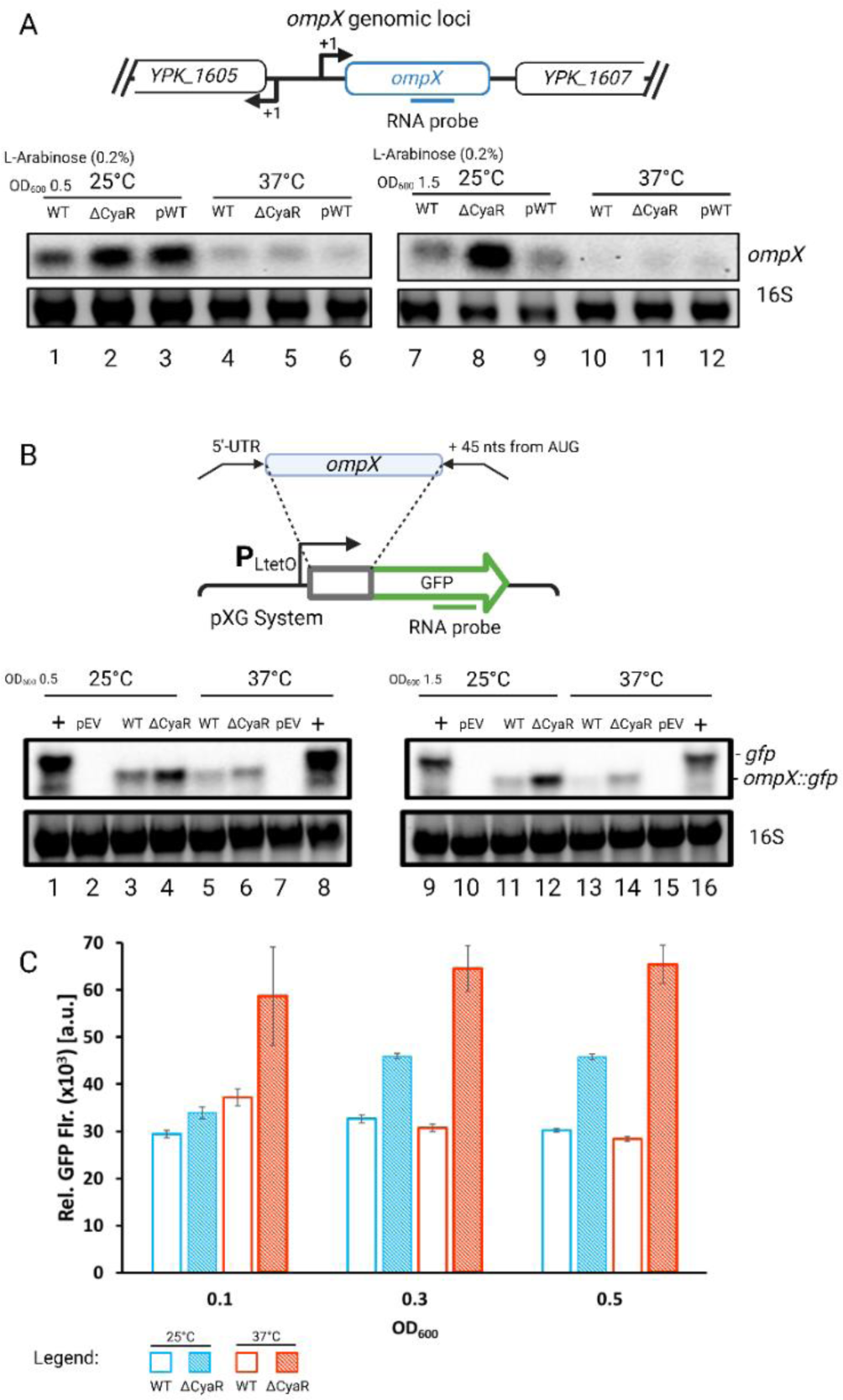
*ompX* is regulated by CyaR in *Y. pseudotuberculosis.* (A) *ompX* mRNA levels were determined by Northern blot analysis of total RNA isolated from early and late exponential cells (OD_600_ of 0.5 & 1.5) of *Y. pseudotuberculosis* WT and ΔCyaR strains carrying the empty vector pGM930, and the ΔCyaR strain carrying pGM930-CyaR. Cultures were grown at 25°C and 37°C and CyaR expression was induced by the addition of arabinose (Ara) for 10 min. (B) Plasmid-derived *ompX::gfp* mRNA was detected by Northern blot analysis of total RNA from early late exponential phase cells (OD_600_ of 0.5 and 1.5) of *Y. pseudotuberculosis* WT carrying the pXG *gfp* vector (+), pXG empty vector (EV), and the WT and CyaR mutant carrying the *ompX::gfp* fusion plasmid. Bacteria were cultivated at 25°C or 37°C. Representative blots of three biological replicates are shown. To ensure equal loading, 10 µg of total RNA was loaded per sample, and gel red stained 16S rRNA served as a loading. (C) GFP fluorescence (flr.) of *Yersinia* WT and ΔCyaR carrying pXG *ompX::gfp* fusions were measured in a plate reader. Relative fluorescence values were calculated as described in Material and Methods.

To isolate the expression of *ompX* from upstream regulation by transcription factors, we constructed a GFP reporter fusion comprising the entire 5’-UTR up to 45 nucleotides into the coding sequence, under the control of a constitutive promoter from the pXG system [38] (Fig. 5B). Northern blot experiments detecting the *gfp* transcript confirmed the role of CyaR as a negative regulator of *ompX* expression Notably, *ompX* transcript levels were consistently higher in the ΔCyaR strain at both temperatures and during both growth phases.

Finally, we took advantage of the expressed *gfp* fusions to measure the fluorescence of the produced GFP protein from bacterial cultures at different optical densities (Fig. 5C). Once more, our findings provided evidence for CyaR being a negative regulator of *ompX* expression. We observed that GFP fluorescence was highest in the absence of CyaR across all tested conditions.

### The *ompX* mRNA is a direct target of CyaR

We employed in-line probing to determine whether *ompX* is a direct target of CyaR and to precisely map the interaction region. This method relies on the principle that the 3’-5’ phosphodiester bond of an unstructured RNA adopts an in-line geometry, rendering it susceptible to nucleophilic phosphor-transesterification by divalent metal ions. This process leads to spontaneous cleavage at the 3’ site of the RNA chain [48, 49].

Subtle structural changes were already noticeable when 5’-[^32^P] labelled *in-vitro* synthesized *ompX* was subjected to in-line structure probing in the absence of CyaR. Nucleotides in the SD and anti-SD regions became susceptible to in-line cleavage at 37°C, indicating that the *ompX* 5’-UTR itself is temperature-responsive (Fig. 6A, lanes 3 and 17). The results obtained from three independent experiments were quantified and normalized to cytosine at position 34, serving as an intramolecular reference, which received a value of 1.0 or 100%. Given its exposed position in the loop between SD and anti-SD region (Fig. 6C), it was efficiently cleaved at both temperatures and unresponsive to CyaR (Fig. 6A).

**Figure 6.**
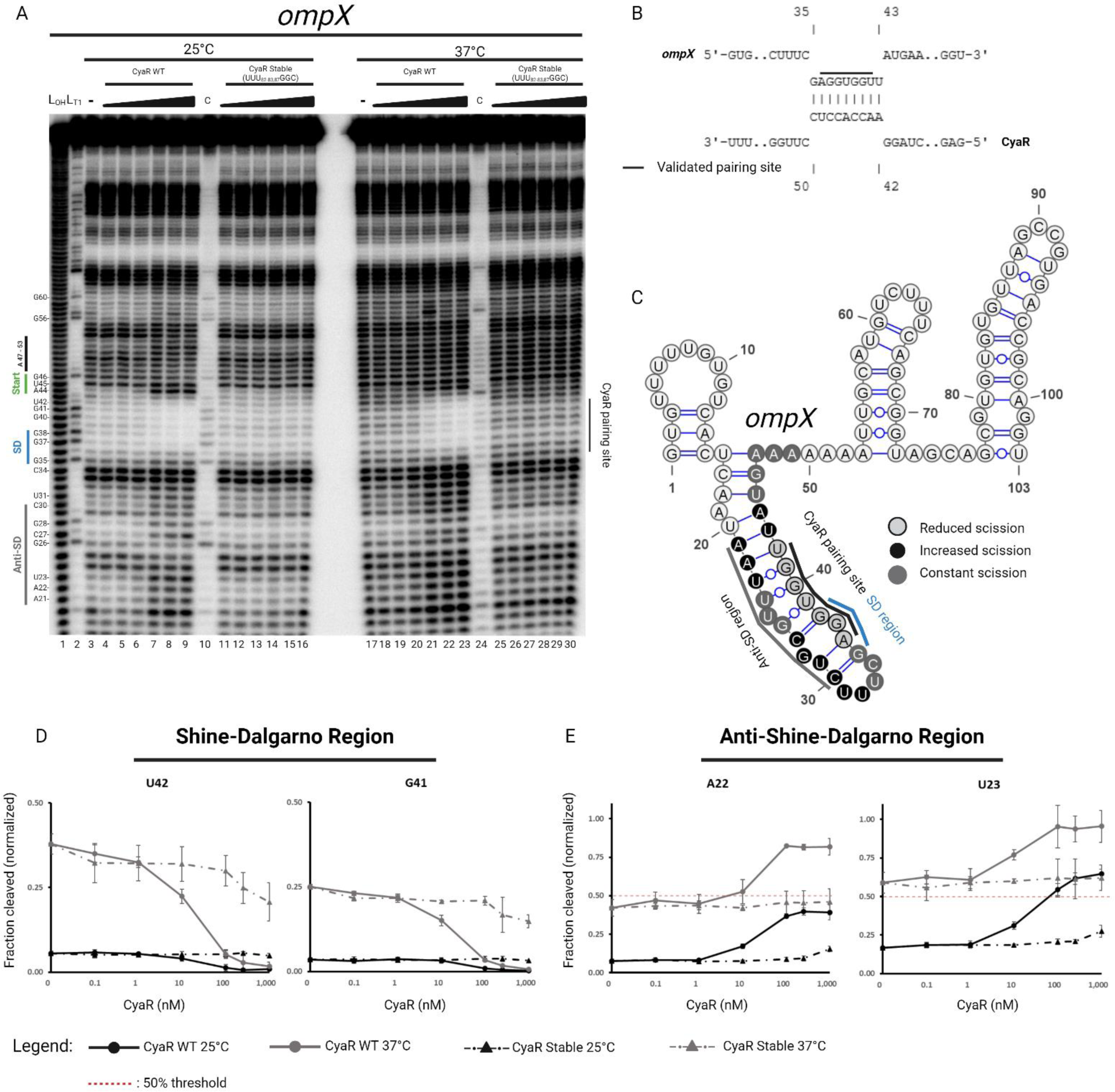
CyaR binds to the SD region of *ompX* and a concerted structural remodeling of *ompX* and CyaR allow for better interaction at 37°C. (A) In-line structure probing of 5’ ^32^P-labeled *ompX* (-43 to +60 from AUG) transcript incubated with MgCl_2_ in the absence (-) or presence of increasing concentrations (0.1 nM, 1 nM, 10 nM, 100 nM, 250 nM, 1000 nM) of CyaR WT or CyaR Stable. Samples were incubated for 40 h at 25°C and 10 h at 37°C. From Left to right: L_OH_: Alkaline ladder; L_T1_: T1 treated RNA at 37°C. (B) Interaction site between *ompX* and CyaR as predicted by the program IntaRNA (C) Secondary structure of the *ompX* 5’-UTR. The SD and anti-SD regions as well as the mapped CyaR binding site are indicated. Nucleotides protected from spontaneous cleavage in the presence of CyaR are in light gray, whereas positions with an increased occurrence of spontaneous cleavage events are shaded black. Constant scission events in and around the stem structure (-23 to +5 from AUG) are indicated in gray. (D, E) Results for individual nucleotides based on densitometric analysis using Image Lab v6.1 (BioRad). Fractions of spontaneous cleavage were normalized to cytosine 34 of the same sample. The normalized fraction of spontaneous cleavage of representative nucleotides (mean ± SD, n=3) from the SD sequence (D) and anti-SD sequence (E). The in-line probing experiment shown above represents one of three independent experiments.

The differences observed in the fragmentation patterns of nucleotides ranging from position 21 to 42 at 25°C compared to 37°C in the absence of CyaR suggest a highly structured stem at 25°C, which becomes liberated at 37°C. The nucleotides surrounding the SD region were spontaneously cleaved at less than 10% at 25°C, contrasting with approximately 25% cleavage at 37°C (Fig. 6A, lanes 3 and 17; quantifications in Fig. 6D and Fig. S4). Based on minimum free energy (MFE) calculations, the hairpin structure spanning residues 16 to 47 is thermodynamically favoured with a value of -6.81 kcal/mol at 25°C, and becomes less stable at 37°C with a value of -3.40 kcal/mol. Most likely, the bulged residue U25, along with the flanking G-U base pairings, contribute to weakening the structural integrity of this stem (Fig. 6C). Indeed, our calculations confirmed that U25 spontaneously cleaves at both 25°C and 37°C, with cleavage rates of 65 ± 4.7% and 84 ± 5.1%, respectively (Fig. S4).

Clear protection against RNA cleavage in the SD region of *ompX* was evident upon the addition of increasing amounts of CyaR at both 25°C and 37°C (∼1% cleavage at 1 µM). Specifically, six nucleotides in *ompX* (37-GGUGGU-42) were shielded by CyaR (43-ACCACC-48), while nucleotide A44 of *ompX* became hypersusceptible to in-line cleavage. Importantly, the mapped interaction region aligns with the *in silico* predicted base-pairing between *ompX* and CyaR (Fig. 6B). Further evidence for an interaction of CyaR with the SD region of *ompX* derives from the concomitant increase of in-line cleavage in the anti-SD region while CyaR protects the SD sequence (Fig. 6A and C). The binding of CyaR to *ompX* should lead to the release of the anti-SD region and a corresponding increase in cleavage events. This was indeed observed at both temperatures, and was more pronounced at 37°C. On average, nucleotides in the anti-SD region displayed spontaneous cleavage rates of approximately 50% at 25°C in the presence of 1 µM CyaR, while cleavage events at the same positions at 37°C occurred at approximately 90% (Fig. 6A, compare lanes 3 – 9 and 17 - 23; quantifications in Fig. 6E and Fig. S4).

Notably, such structural rearrangements in *ompX* were not observed when a stable, temperature unresponsive CyaR variant, in which the U residues 82, 83 and 87 were changed to G, G and C, respectively (Fig. 2C), was added. At 25°C, there was less than 1% difference in the absence of CyaR or with 1 µM of CyaR Stable (Fig. 6A, lanes 11 – 16). However, at 37°C, a slight decrease of about 10% in spontaneous cleavage in the SD region was observed from 0 nM to 1 µM of CyaR Stable (Fig. 6A, lanes 25 – 30; quantifications in Fig. 6D and Fig. S4). These findings, together with the probing results in Fig. 2A, suggest that the CyaR Stable population does not exhibit homogenous closure in structure, as a subpopulation can bind to *ompX* at 37°C.

### The 5’-UTR of *ompX* itself is thermo-responsive

The in-line probing experiments suggested that the *ompX* mRNA has RNA thermometer-like properties, as it released the SD sequence at increasing temperature (Fig. 6). To provide further evidence for this dynamic behaviour, we took advantage of our previous PARS analysis [25]. The drop in PARS profile in both the SD region and the complementary anti-SD region at 37°C (Fig. 7A) indicates that this region becomes single-stranded with increasing temperature. Additional support for RNA thermometer-like translational control was obtained by a reporter gene fusion of the *ompX* 5’-UTR to *bgaB* coding for a thermostable β-galactosidase. The well-established thermometer upstream of *yopN* [28] was used as a positive control. In addition to the WT *ompX* fusion, two designed variants presumed to be stabilized and repressed (R1 and R2; Fig. 7B) were examined. In contrast to the 7-fold increase in β-galactosidase activity observed by the *yopN* fusion, the *ompX* fusion displayed only a modest 2-fold increase in β-galactosidase at 37°C compared to 25°C (Fig. 7C). The mutations introduced in R1 (UU_23-24_CC) were not sufficient to suppress translation, necessitating an additional mutation in R2 (G_28_C) to completely abolish β-galactosidase activity at both temperatures (Fig. 7C).

**Figure 7.**
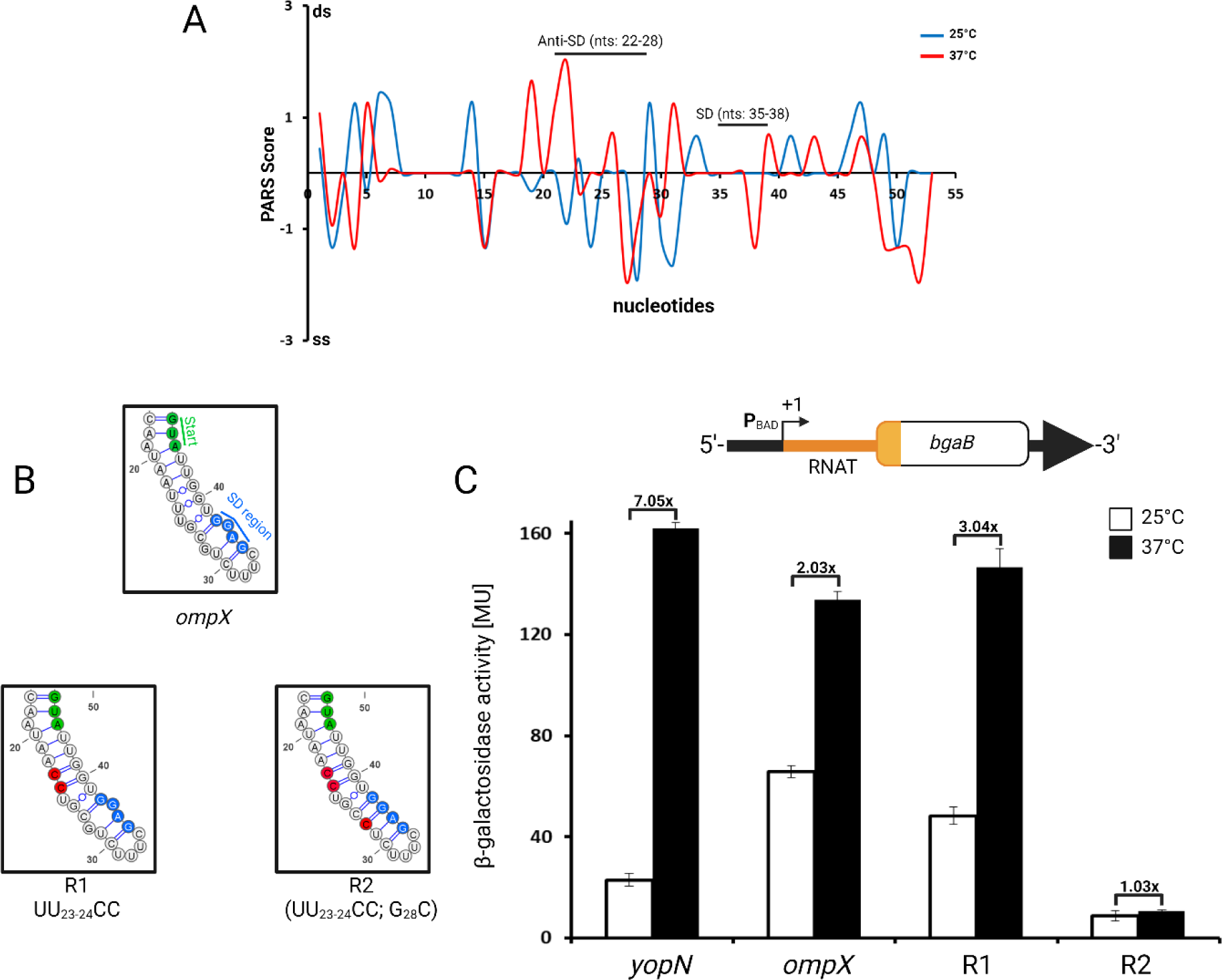
The *ompX* 5’-UTR contains a weak RNA thermometer. (A) PARS profile of the *ompX* 5’-UTR (B) Nucleotides in blue and green represent the SD sequence and start codon, respectively. The boxes show the secondary structures from nucleotides 17 to 46 of the WT and mutated R1 and R2 sequences. Nucleotide exchanges are depicted in red. (C) Reporter gene assay of *yopN-* (control) and *ompX*-bgaB fusions. *Y. pseudotuberculosis* YPIII carrying pBAD2 fusion constructs with the indicated fusions was grown to an OD_600_ of 0.5. Transcription of the *bgaB* fusions was then induced by L-arabinose (0.1% (w/v)). The cultures were split with half of the main culture left at 25°C and the other half transferred into flasks prewarmed at 37°C. The cultures were incubated for further 30 minutes prior to β-galactosidase activity measurements. Shown are the mean activities (n= at least 3 biological replicates) with standard deviations.

The results (Fig. 6 and 7) collectively suggest that the *ompX* 5’-UTR exhibits weak RNA thermometer-like properties. Based on the combined structure probing results, we conclude that both partner RNAs, the mRNA *ompX* and the sRNA CyaR are temperature responsive. The demonstrated binding of CyaR to the SD region of *ompX* (Fig. 6) suggests that the sRNA can modulate translation initiation from the *ompX* mRNA. This hypothesis was tested in the next experiment.

### CyaR represses translation initiation better at 37°C

To demonstrate the effect on translation through the interplay of both transcripts, we employed primer extension inhibition assays (toeprinting). Transcripts were synthesized *in vitro*, refolded and incubated with the 30S subunit, or both CyaR WT transcript and 30S, at different temperatures prior to reverse transcription. The successful formation of a 30S translation initiation complex can be visualized by the presence of a truncated cDNA product, called toeprint. A clear toeprint signal was observed in the presence of 30S ribosomes at the appropriate position 15 to 17 residues upstream of the first nucleotide of the start codon (Fig. 8). Consistent with the observed RNAT-like opening of the *ompX* secondary structure, (Fig. 6 and 7), this signal increased 2.5 and 3.2-fold at 37 and 42°C, respectively. In the presence of CyaR, the toeprint signal was reduced to 75 and 55% at 25 and 37°C (Fig. 8) indicating that the sRNA interfered with translation initiation. At 42°C, the toeprint was comparable to the one in the absence of CyaR, suggesting that the mRNA/sRNA complex becomes unstable and/or the 30S subunit is more competent in binding to the SD region of *ompX* than CyaR under these conditions.

**Figure 8.**
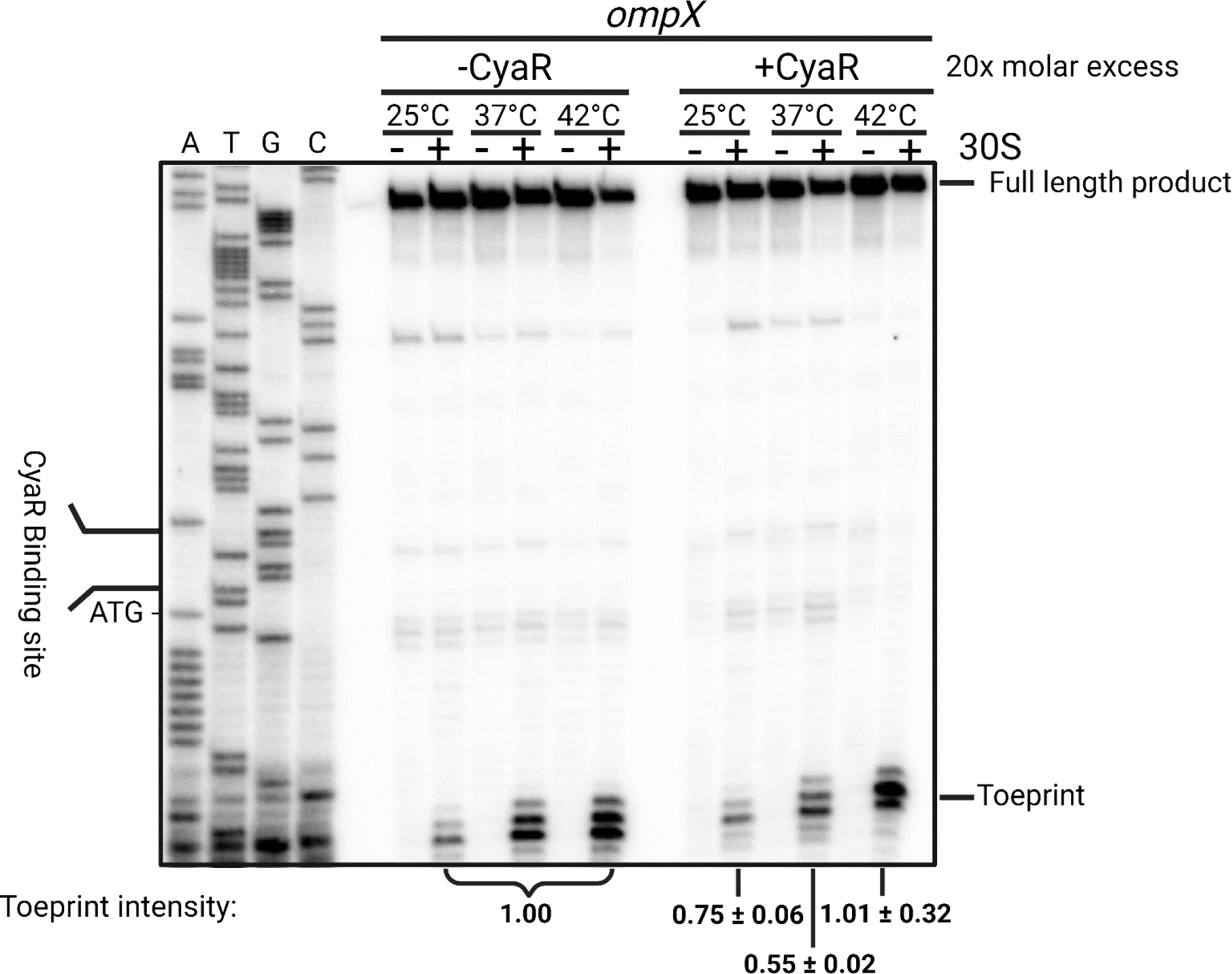
Translation initiation is reduced at 37°C when CyaR is present *in vitro.* Primer extension inhibition assay of the *ompX* transcript at 25, 37, and 42°C in the presence (+) or absence (-) of 6 pmols of 30S ribosomal subunit and the presence (+CyaR) or absence (-CyaR) of 1.5 pmol CyaR WT. A DNA sequencing ladder (ATGC) of *ompX* serves as orientation. Full length product and a characteristic toeprint signal (+15-17 nt from the A of AUG) due to the bound ribosome is indicated. Toeprint intensities were quantified by Image Lab v6.1 (BioRad). The signals at different temperatures without CyaR were set to 1, and relative changes were determined for the samples with CyaR. The experiment is a representative of two independent replicates.

### *ompX* mRNA decay is accelerated by CyaR at 37°C

Aside from translational repression, sRNAs can influence the turnover rates of their regulated transcripts [50, 51]. To assess the effect of CyaR on the stability of the *ompX* mRNA, we measured the half-lives of both RNAs using Northern blot analyses. Bacteria were cultured at 25°C, after which half of the culture was shifted to 37°C before transcription was stopped by the addition of rifampicin. CyaR was found to be highly stable at both temperatures (Fig. 9A). In contrast, the half-life of *ompX* varied with temperature and the presence of CyaR. Most notably, the stability of the *ompX* transcript in the WT strain was reduced from 7.4 min at 25°C to 1.6 min at 37°C (Fig. 9B). In the ΔCyaR strain, the half-life of *ompX* increased to more than 10 min at 25°C and to 2.6 min at 37°C min, suggesting that CyaR accelerates the degradation of the *ompX* transcript.

**Figure 9.**
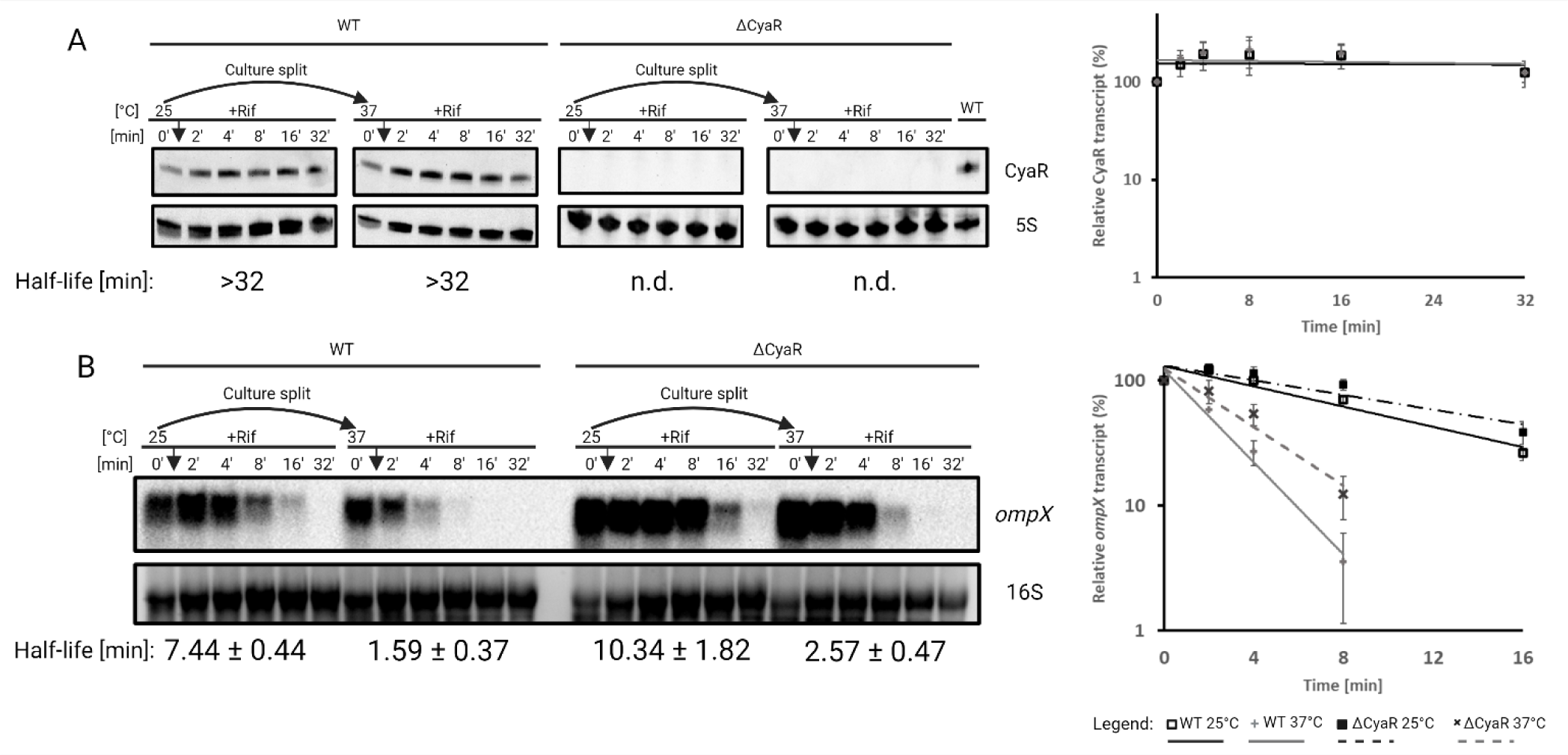
*In vivo* turnover of *ompX* is accelerated at 37°C when CyaR is present. Northern blot analysis of *in vivo* RNA decay of *ompX* and CyaR of *Y. pseudotuberculosis* WT and ΔCyaR strains after rifampicin treatment. Exponentially growing cells (OD_600_ of 0.5) grown at 25°C were split into flasks at 25°C and prewarmed flasks at 37°C. Samples were collected prior to addition of rifampicin (0’), then at the indicated timepoints (2’ – 32’). (A) 10 µg of total RNA were loaded per sample and probed for CyaR. A WT sample was added to both 25 and 37°C blots of ΔCyaR as additional control, which was cropped out of the 25°C blot for symmetry. Graphical representation of *in vivo* CyaR decay (mean ± SD, n=3) computed based on the Northern blot bands. The band before addition of rifampicin was set to 100%. (B) 10 µg of RNA from the same experiment above was loaded per sample and probed for *ompX*. Gel-red stained 16S rRNA serves as loading control. Half-lives shown below the blots are based on decay rates calculated from the graph shown to the right of the respective blots (mean ± SD, n=3 biological replicates). Trendlines were fitted to a first order exponential decay function.

### OmpX protein is preferentially produced at low temperature

Given the preferential interaction between CyaR and *ompX* at 37°C due to conformational rearrangements, one might expect higher levels of OmpX protein at lower than at higher temperatures. To test this hypothesis, we integrated an *ompX*^3xFLAG^ gene encoding a C-terminally FLAG-tagged OmpX protein, into the chromosome. We observed a significant reduction in OmpX levels at 37°C compared to 10, 20, and 25°C in exponentially growing cells (Fig. 10A). Shifting cultures grown at 25°C to 37°C or 10°C resulted in reduced OmpX levels when the temperature was upshifted (Fig. 10B). Similarly, upshifting from 10°C to higher temperatures led to decreased OmpX levels, especially at 37°C (Fig. 10C) while downshifting from 37°C to 25°C restored higher OmpX levels (Fig. 10D). Measuring OmpX-FLAG levels in various mutants revealed that Hfq plays a role in regulation, whereas OmrA, another sRNA, does not. Consistent with a temperature-modulated negative role of CyaR on *ompX* expression, deletion of the sRNA resulted in an ∼40% increase in OmpX levels at 25°C and a twofold increase at 37°C compared to the WT (Fig. 10E).

**Figure 10.**
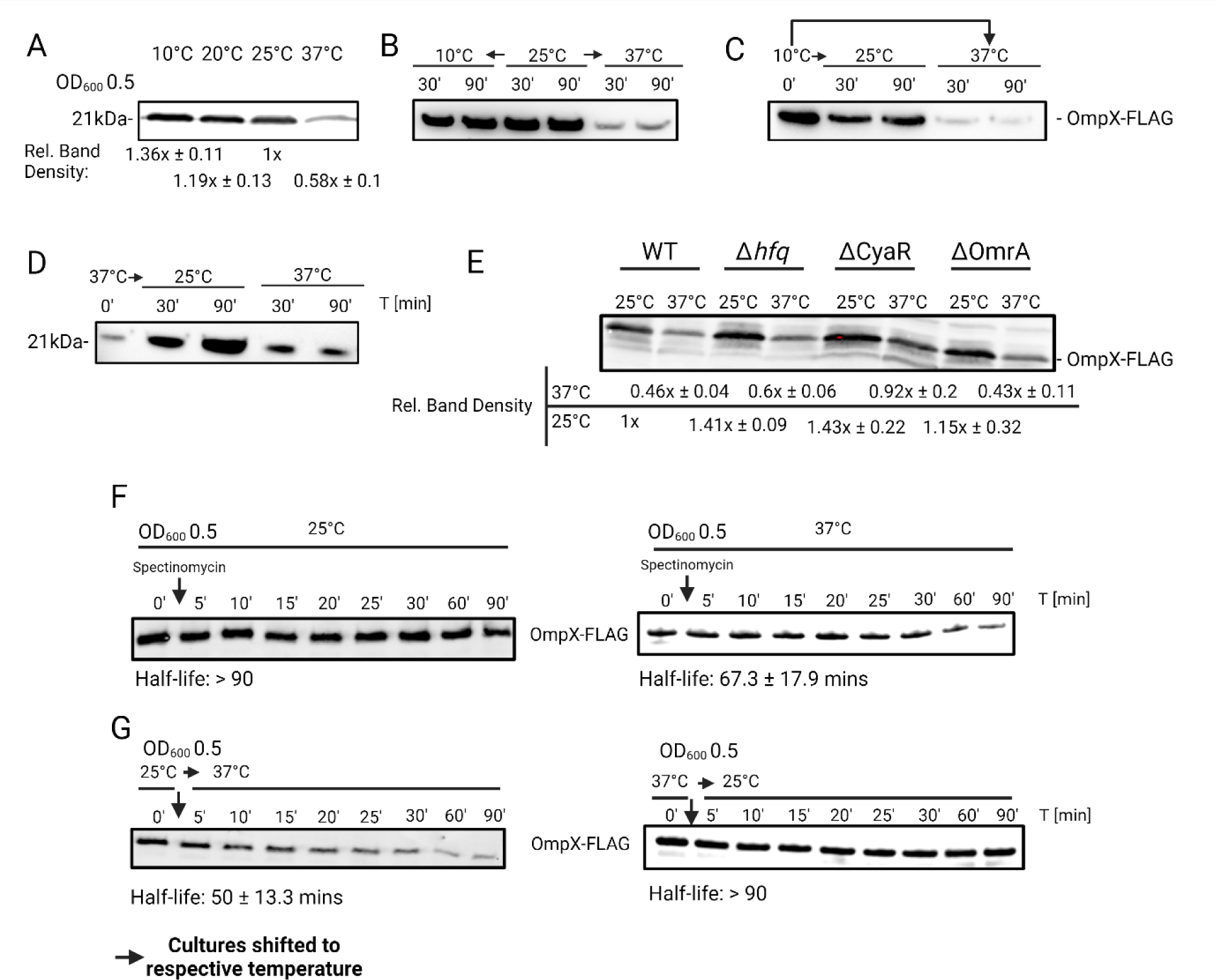
OmpX protein levels are highest at 25°C. (A-D) Western blot analysis of 3xFLAG-tagged OmpX protein levels harvested from exponentially growing cells (OD_600_ of 0.5) at indicated temperatures. (B-D) Samples were harvested prior to shifting at the indicated temperature then harvested again after shifting to the indicated temperature at the indicated time point. Protein amounts were adjusted to an OD_600_ of 0.5. To ensure equal loading of samples, nitrocellulose membranes were stained with Ponceau S. (E) Western blot analysis of 3xFLAG-tagged OmpX protein levels harvested from exponentially growing cells (OD_600_ of 0.5) of *Y. pst* WT, Δ*hfq,* ΔCyaR, and ΔOmrA at indicated temperatures. To ensure equal loading of samples, TGX stain-free gels (BioRad) were activated under UV-light for 45 secs prior to transfer onto a nitrocellulose membrane. (F & G) Western blot analysis of *in vivo* protein degradation of 3xFLAG-tagged OmpX protein levels harvested from exponentially growing cells (OD_600_ of 0.5) of *Y. pst* WT at the indicated temperature. Samples were harvested prior to addition of spectinomycin (0’), then at indicated timepoints (5’ – 90’). (F) Cells were initially grown at the indicated temperature to the optical density mentioned above, then shifted up (G; left panel) or down (G; right panel) (indicated by arrows). Half-lives below the blots are based on decay rates calculated from the graph in (Supplemental Figure 5). Protein amounts were adjusted to an OD_600_ of 0.5. To ensure equal loading of samples, TGX stain-free gels (BioRad) were activated under UV-light for 45 secs prior to transfer onto a nitrocellulose membrane. Protein amounts were adjusted to an OD_600_ of 0.5. For quantitative western blots, TGX stain-free gels (BioRad) were activated under UV-light for 45 secs prior to transfer onto a nitrocellulose membrane. Densitometry measurements represent mean ± SD, n= minimum of 3 biological replicates.

To investigate whether the temperature-dependent OmpX levels were caused by changes in protein stability, we determined the half-lives of OmpX after halting protein biosynthesis with spectinomycin. While OmpX was a stable protein at 25°C, it was prone to slow degradation at 37°C (Fig. 10F). Further evidence for a degradation of OmpX at elevated temperatures was derived from shift experiments. When cultures were initially grown at 25°C and then shifted to 37°C, OmpX displayed a half-life of approximately 50 min (Fig. 10G). However, the protein remained stable when cultures were shift from 37 to 25°C.

In summary, our results show that the expression of *ompX* is effectively controlled by numerous post-transcriptional mechanisms controlling *ompX* mRNA stability and translation as well as OmpX protein stability.

## DISCUSSION

### The regulatory potential of environmentally responsive mRNA structures

Structured RNA regions are ideally suited to monitor environmental conditions. They can either detect chemical cues through specific ligand binding or perceive physical parameters, such as temperature fluctuations, by structural rearrangements. Despite two decades of riboswitch discovery, which has led to the identification of over 55 classes of distinct classes of riboswitches, it is believed that this only represents a small fraction of the riboswitches existing in nature [52]. Recently, a bioinformatic pipeline predicted 44 novel riboswitch candidates highlighting the vast structural and functional diversity in regulatory RNA motifs [53]. Moreover, the biochemical repertoire of ligands sensed by validated riboswitches is extremely broad and ranges from simple ions to large and complex organic compounds such as nucleotides, amino acids or enzyme cofactors [54].

While riboswitches typically rely on some degree of sequence and structure conservation for ligand binding, facilitating their identification through *in silico* searches, RNATs are more challenging to identify by bioinformatic pipelines. Their temperature-labile RNA structures, which block ribosome binding at low temperatures, do not necessarily require a high degree of sequence conservation. Nevertheless, some bioinformatic strategies utilizing previously identified RNATs as templates have been successful [55, 56]. Experimental transcriptome-wide approaches have uncovered numerous structurally diverse RNAT candidates by resolving RNA structures at different temperatures. As current champion, *Y. pseudotuberculosis* has been found to harbour dozens of temperature-responsive RNA structures upstream of not only virulence genes but also heat shock and metabolic genes [25, 26]. Interestingly, the number of RNAT candidates not associated with apparent temperature-dependent processes, such as the heat shock or virulence response, is on the rise. Transcriptome-wide RNA structuromics in *Bacillus subtilis* for example, has revealed RNATs governing the expression of glycerol permease genes [57]. Additionally, a fourU-type RNAT has been identified in the 5’-UTR of *blyA* of the *B. subtilis* phage SOβc2 [58]. The *blyA* gene encodes an autolysin involved in cell wall hydrolysis during the lytic life cycle of the phage. Other RNATs have been described upstream of gene encoding a σ70 RNA polymerase subunit in *Mediterraneibacter gnavus* and *Bacteroides pectinophilus*, as well as in the bacteriophage Caudoviricetes, which infects *B. pectinophilus* [56]. Yet another RNAT resides in the 5’-UTR of the tetracycline resistance gene *tetR* in *E. coli* and *Shigella flexneri*. While these findings significantly advance our understanding of the prevalence of RNATs in bacterial transcriptomes, they raise the question of why all these genes are temperature-regulated and how other environmental cues might be integrated into regulation.

### A thermo-responsive small RNA in *Y. pseudotuberculosis*

Our study expands the scope of thermo-responsive RNA structures by presenting evidence for the existence of a functionally relevant RNAT-like structure in the sRNA CyaR. CyaR homologs from closely related enteric species such as *E. coli* and *Salmonella* exhibit divergent expression patterns, distinct sequences (Fig. 1A), and predicted structures to that of *Y. pseudotuberculosis* and *Y. pestis* [59]. In *E. coli* [33] and *Salmonella* [13], CyaR accumulates during transition to stationary phase. Notably, in *Y. pseudotuberculosis*, significant differences in CyaR expression were observed in low-density cultures, with markedly higher CyaR levels at 37°C compared to 25°C (Fig. 3A-C; Nuss et al., 2015), suggesting a temperature-dependent regulatory function.

It is remarkable that the seed region of CyaR undergoes a temperature-responsive structural transition that facilitates interaction with the *ompX* mRNA, which has RNAT-like properties itself (Fig.11A). The seed region of sRNAs is integral to their function, and exchanging the seed region of one sRNA to another one changes its regulatory function [36]. As the critical ‘regulatory module’ of any sRNA, the seed region should be structurally available for pairing with its target mRNA and not be locked in a stem-loop [60]. Structural evidence for CyaR in *Y. pseudotuberculosis* shows that its seed region is paired with its 3’-polyU tail at ambient temperatures (Fig. 2A). The hidden seed region at 25°C restricts its pairing efficiency with its target *ompX* (Fig. 6). The cumulative structural evidence suggests that both in the CyaR and *ompX* transcripts intramolecular folding is favoured at 25°C. As the temperature increases, both structures open to facilitate intermolecular interactions (Fig. 11A).

**Figure 11.**
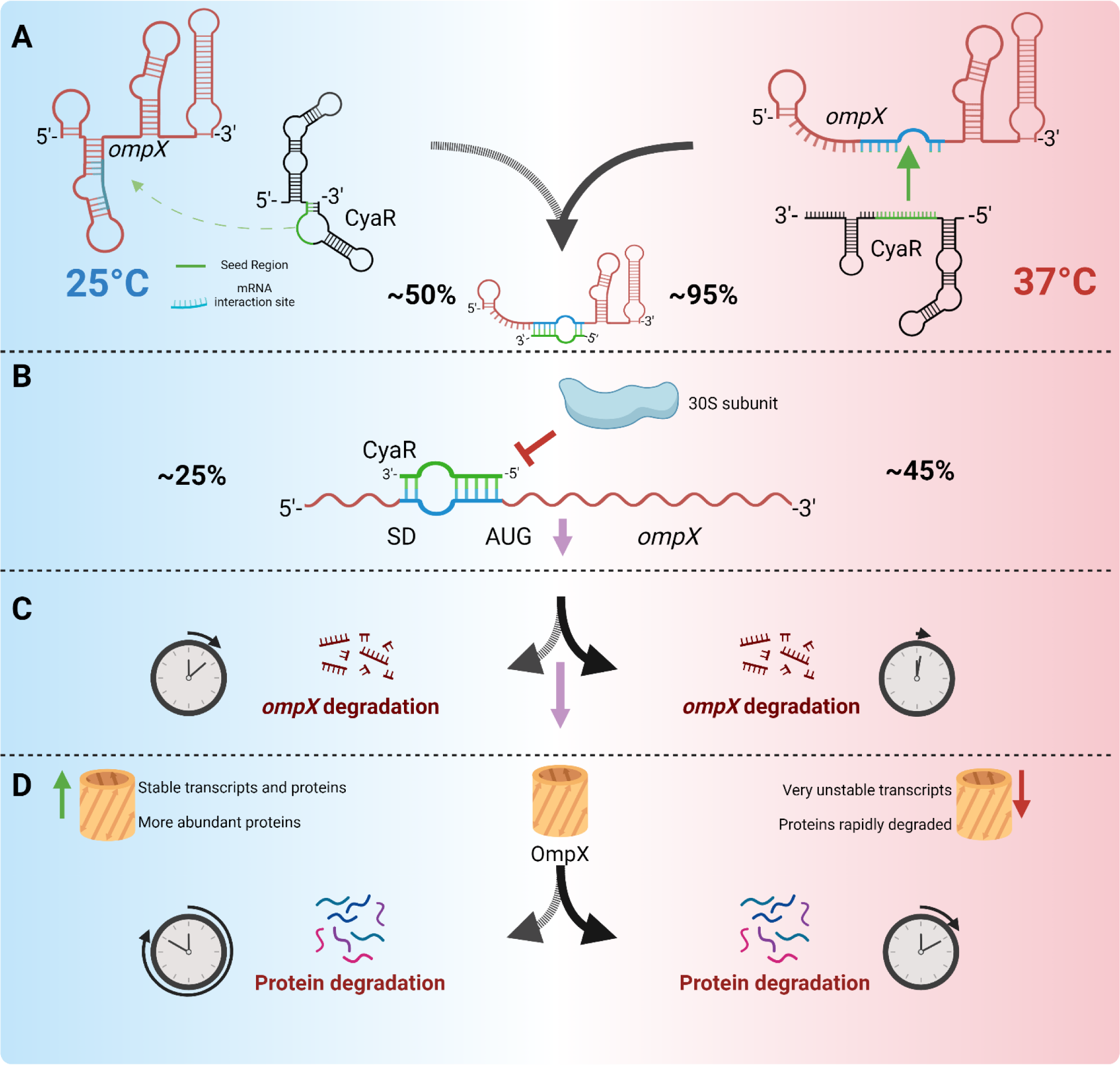
Mechanistic overview of temperature-dependent post-transcriptional – translational regulation of *ompX*. (see text for details).

Base pairing between CyaR and the *ompX* 5’-UTR has consequences on both translation initiation and mRNA stability. As shown *in vitro* by toeprinting assays, the facilitated interaction between CyaR and *ompX* at 37°C reduced ribosome binding by 45% compared to a 25% reduction by CyaR at 25°C (Fig. 11B). Reduced ribosome binding can promote the destabilization of transcripts [61] and we show *in vivo* that the CyaR-dependent reduction of ribosome binding at 37°C stimulated degradation of the *ompX* transcript (Fig. 11C).

These cumulative post-transcriptional events could already be sufficient to explain the reduction of OmpX protein levels at 37°C compared to 25 or 10°C (Fig. 10). However, we find that in addition the stability of OmpX protein is modulated by temperature such that its degradation is accelerated at 37°C (Fig. 11D). We do not know the identity of the protease responsible for OmpX degradation. One potential candidate is the periplasmic protease DegP (HtrA), which is able to access and degrade proteins in the outer membrane [62]

### Temperature-regulated synthesis of outer membrane proteins in *Y. pseudotuberculosis*

The outer membrane acts as the first line of defence in Gram-negative bacteria in an ever-changing environment and requires a constant remodeling to enable the intake of nutrients, to catalyse chemical reactions, and to allow the bacteria to adhere to surfaces or each other. Research in the past decades has established extensive regulatory sRNA networks involved in outer membrane protein biogenesis [63, 64]. Apart from porin-like proteins, sRNA-mediated regulation extends to other outer membrane structures such as multidrug efflux pumps [65], or a haem receptor in enterohemorrhagic *E. coli* that is controlled by CyaR and an RNAT in the *chuA* mRNA [22].

*Y. pseudotuberculosis* is known to remodel most of its outer membrane structures in response to a temperature upshift. Flagellar structures for motility are replaced by virulence factors (e.g. adhesins, invasins) [66] or complex T3SS structures [29, 67]. The outer membrane protein OmpA has been shown to be upregulated by an RNAT at 37°C [30]. This is also true for OmpA in *S. dysenteriae* where it is important for pathogenesis [68]. In contrast, OmpX is ultimately downregulated at 37°C by a cascade of regulatory events (Fig. 11). This is different in *Y. pestis* where OmpX expression is constitutive [69].

The biological function of OmpX is currently unknown. The protein displays extensive homology to Ail as it exists in *Y. pestis, Y. pseudotuberculosis* and *Y. enterocolitica* [69, 70]. In the latter organism, Ail is induced at 37°C [71, 72]. According to our RNA-seq results, transcription of *ailA* is upregulated 2.8-fold at 37°C in *Y. pseudotuberculosis* (Supplementary dataset 2). Moreover, the *ailA* 5’-UTR carries a functional RNAT contributing to upregulation at host body temperature [25]. Comparison of *Y. pseudotuberculosis* Ail and OmpX sequences (Fig. S6A) to *E. coli* OmpX [73] reveals many similarities. Excluding the periplasmic signal sequence, membrane-spanning residues from both sequences are highly conserved (up to 60% sequence identity), whereas most cell surface residues are different (approx. 15% conservation). The OmpX and Ail structures predicted by Alphafold 3 [74] appear highly similar (Fig. S6C; lateral and bottom view). However, the four extracellular loops are distinctive in both proteins (Fig. S6C, top view). In *Y. pestis, ailA* codes for an adhesin, which binds to and neutralizes neutrophils via T3SS-mediated translocation of Yops (*Yersinia* outer proteins) [76, 77]. Positively charged residues on the extracellular surface of Ail were hypothesized to be involved in adhesion to host cells and promote Yop delivery into phagocytic and epithelial cells [78, 79]. We hypothesize that the *ailA* transcript with its RNAT [25] lies in wait, ready for translation as *Y. pseudotuberculosis* encounters a temperature upshift. This raises the intriguing question of whether OmpX is the cold-induced counterpart of Ail and what physiological effects are associated with it.

## DATA AVAILABILITY

Data is available at the Gene Expression Omnibus (GEO) database under the accession number GSE270081 (https://www.ncbi.nlm.nih.gov/geo/query/acc.cgi?acc=GSE270081).

## SUPPLEMENTARY DATA

Supplementary Data are available at NAR online.

## ACKNOWLEDGEMENTS

We are grateful to Petra Dersch for providing the pYP32 plasmid used to generate the CyaR deletion mutant and for the *Y. pseudotuberculosis* Δ*hfq*, Δ*crp*, and Δ*csrA* strains used in this study. We thank the RNA group for continuous and fruitful discussion during the writing of this manuscript.

