## supplement for "Two temperature-responsive RNAs act in concert: The small RNA CyaR and the mRNA *ompX*"

### Supplementary Figures

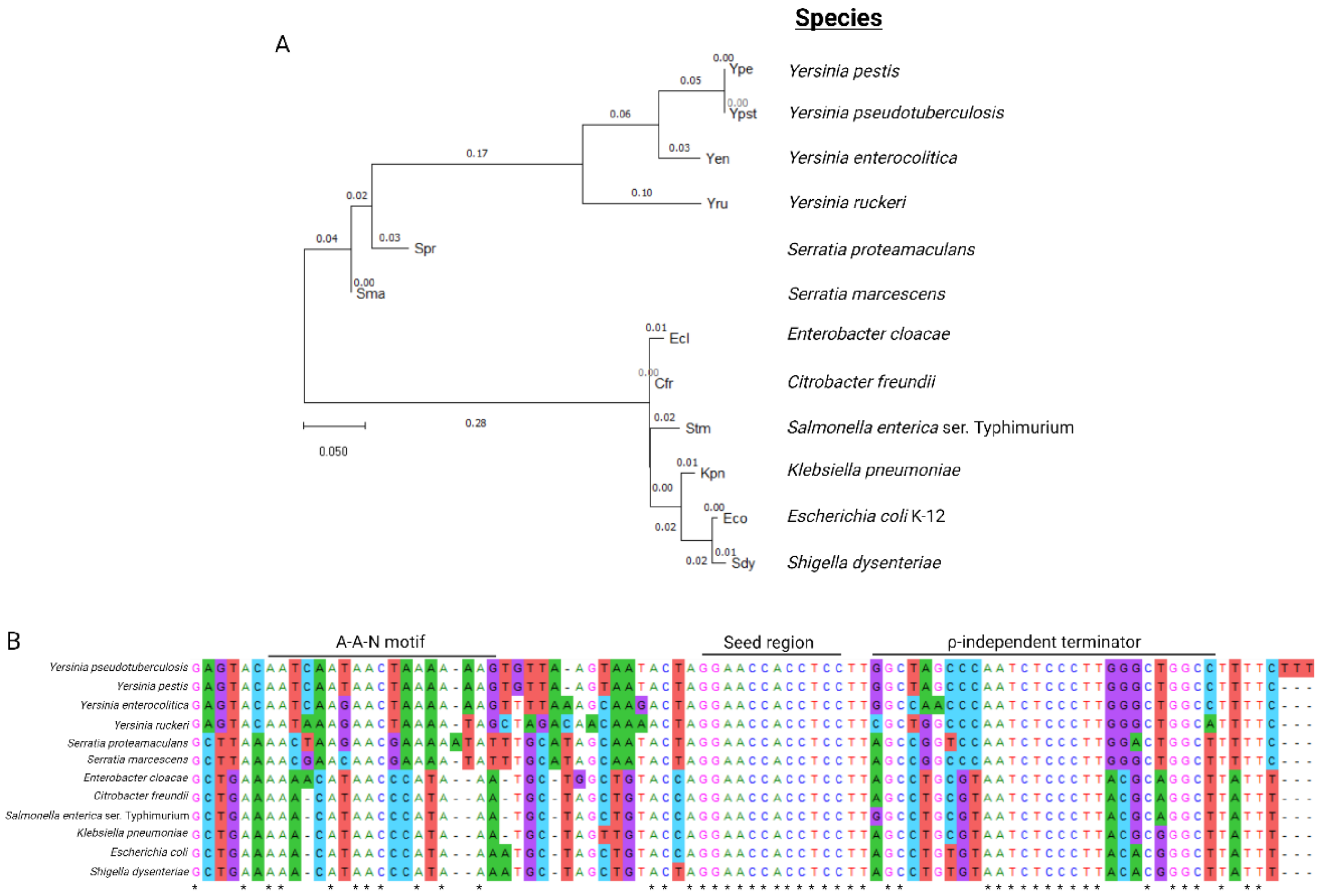

**Supplementary Figure 1. Phylogenetic analysis of CyaR sequences.**

(A) Evolutionary analysis by Maximum Likelihood method. The evolutionary history was inferred by using the Maximum Likelihood method and Tamura-Nei model [1]. The tree with the highest log likelihood (-413.05) is shown. Initial tree(s) for the heuristic search were obtained automatically by applying Neighbor-Join and BioNJ algorithms to a matrix of pairwise distances estimated using the Tamura-Nei model, and then selecting the topology with superior log likelihood value. The tree is drawn to scale, with branch lengths measured in the number of substitutions per site (next to the branches). This analysis involved 12 nucleotide sequences. There was a total of 93 positions in the final dataset. Evolutionary analyses were conducted in MEGA11 [2]. (B) Sequence alignment of 12 representatives of Enterobacteriaceae family. Perfectly conserved nucleotides are denoted with *. Accession numbers: CBWP010000021.1 *Citrobacter freundii*, AXOM01000023.1 *Enterobacter cloacae*, U00096.3 *Escherichia coli* K-12, CP000647.1 *Klebsiella pneumoniae*, AE006468.2 *Salmonella enterica* ser. Typhimurium, HG326223.1 *Serratia marcescens*, CP000826.1 *Serratia proteamaculans*, CP000034.1 *Shigella dysenteriae*, AM286415.1 *Yersinia enterocolitica*, AL590842.1 *Yersinia pestis*, CP009792.1 *Yersinia pseudotuberculosis*, JPFO01000006.1 *Yersinia ruckeri*.

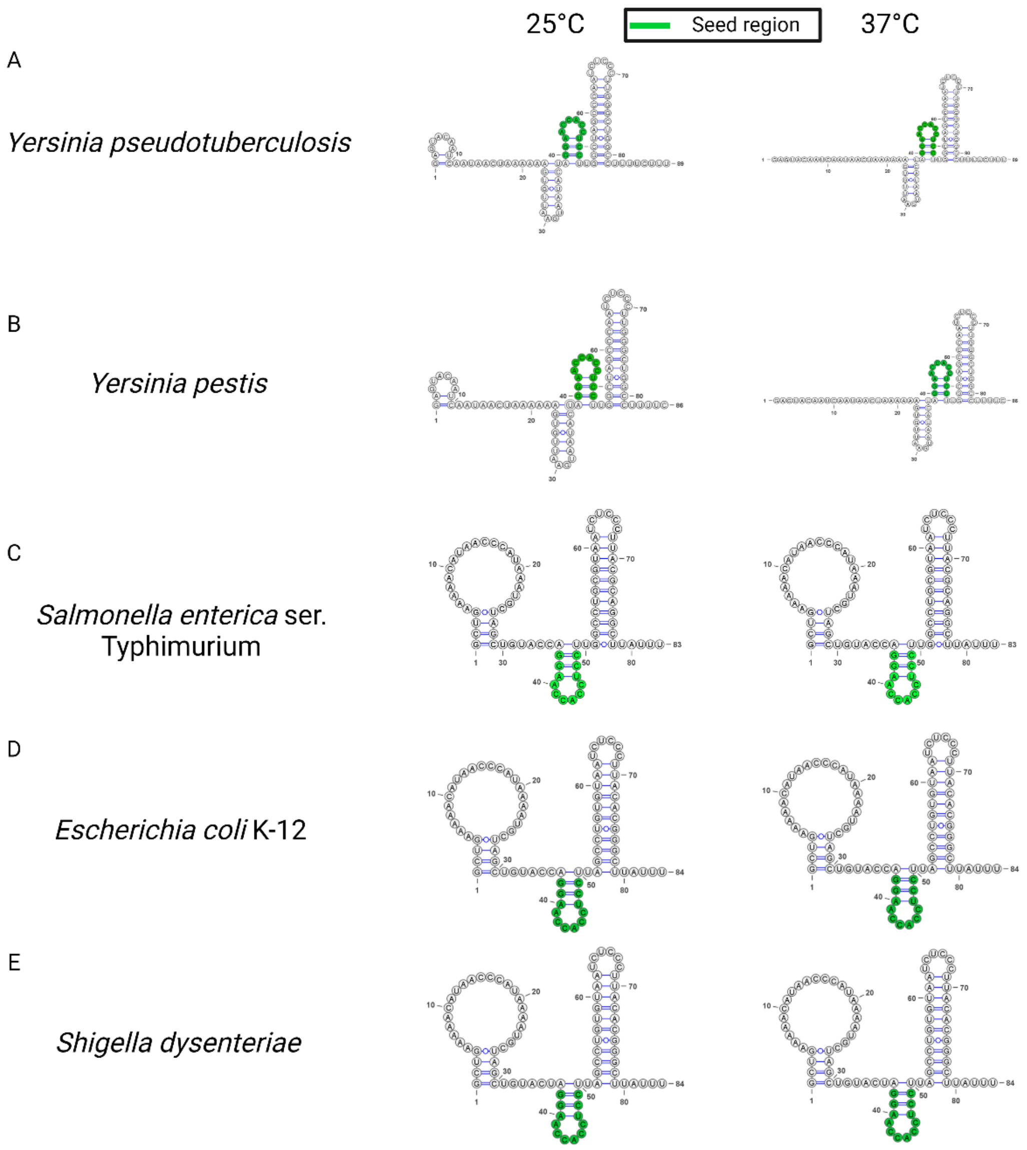

**Supplementary Figure 2. Predicted secondary structures of CyaR.**

Calculated secondary structure of CyaR from representative enterobacterial species via RNAfold webserver (Uni Vienna). Sequences are gathered from the same accession numbers in Fig. S1. Energy parameters were based on Turner model, 2004 and rescaled to 25 or 37°C to guide the algorithm to the most likely base-pairing probabilities.

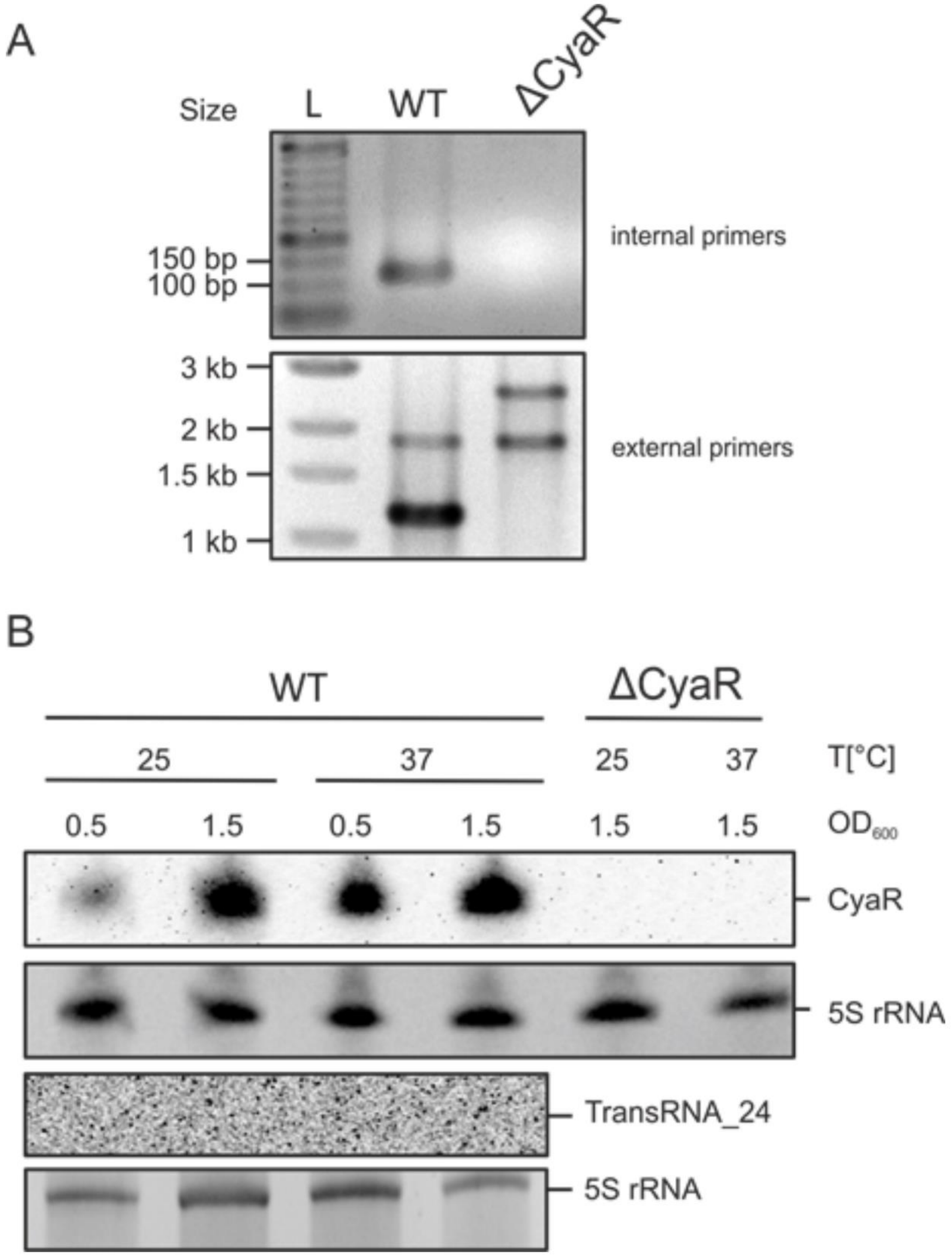

**Supplementary Figure 3. Confirmation of CyaR deletion by PCR and Northern blot.** (A) Replacing CyaR with a kanamycin resistance cassette resulted in changes in DNA fragment sizes using primers that bind within and up- and downstream of the CyaR locus. No DNA fragment was produced using internal primers compared to the wild type (WT). L: DNA ladder. (B) Northern blots confirming the non-expressed transRNA_24 on the complementary strand of CyaR and the deletion of CyaR. A total of 10 µg of RNA was loaded per sample, and 5S rRNA stained with ethidium bromide served as loading control.

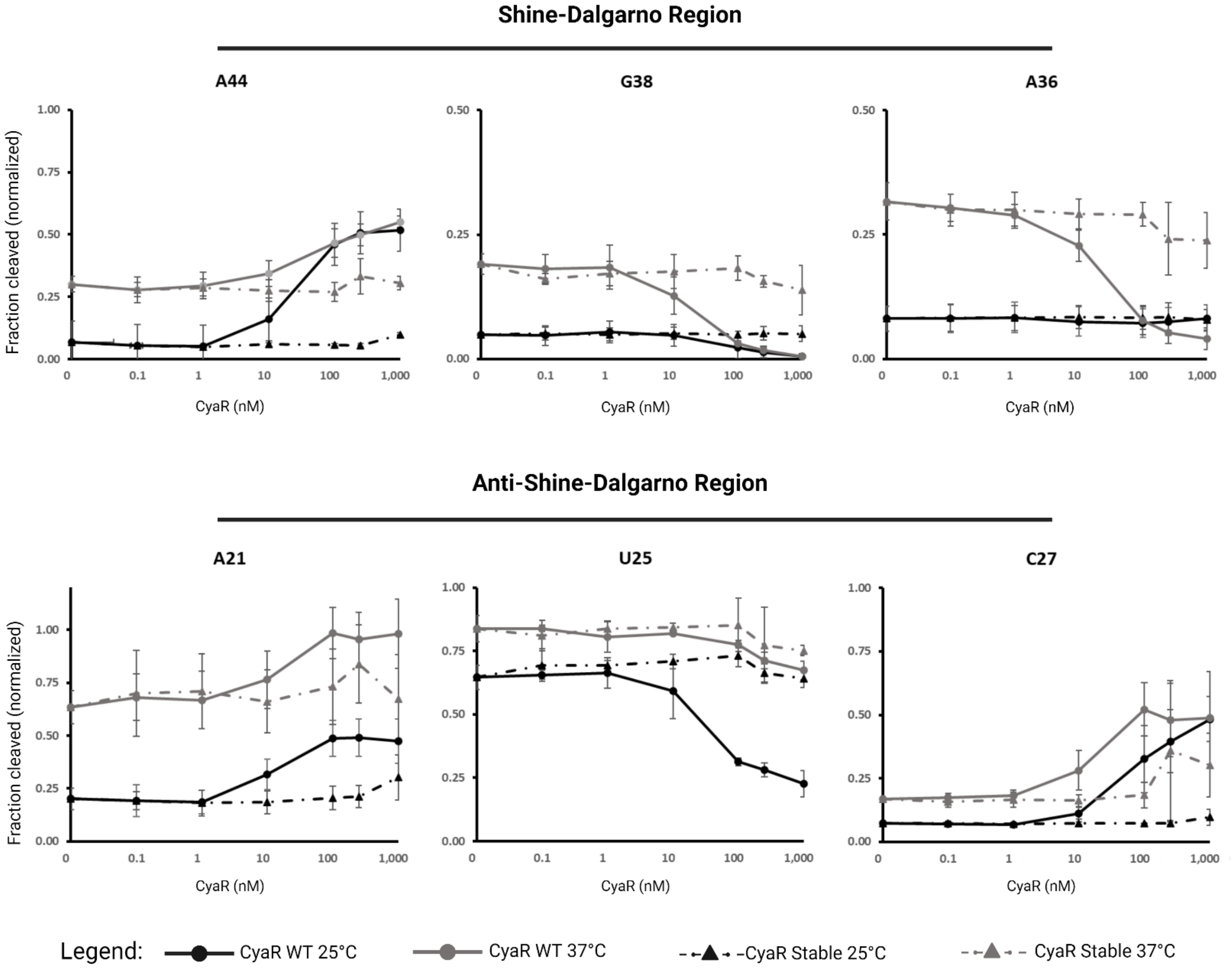

**Supplementary Figure 4. Calculated in-line cleavage of representative nucleotides from the SD region and anti-SD region of *ompX*.**

Densitometry measurement obtained with Image Lab v6.1 (BioRad). Fractions of spontaneous cleavage were normalized to cytosine 34 of the same sample. The normalized fraction of spontaneous cleavage of nucleotides (mean ± SD, n=3).

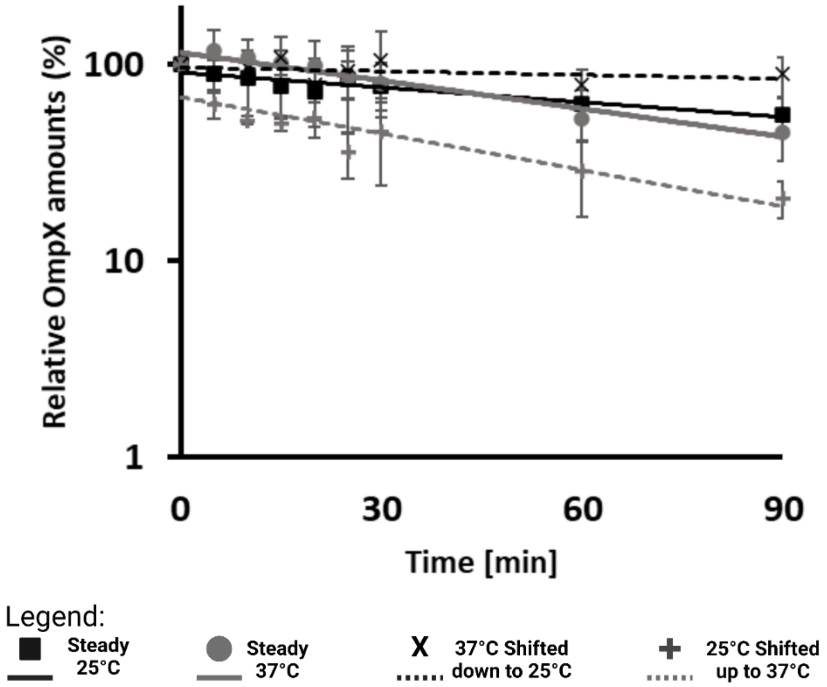

**Supplementary Figure 5. Graphical representation of *in vivo* OmpX-3xFLAG stability experiments**

Densitometry measurements obtained with Image Lab v6.1 (BioRad). Graphical representation of *in vivo* OmpX-3FLAG degradation mean ± SD, n=6 biological replicates for cultures grown steadily at their respective temperatures and n=3 biological replicates for cultures shifted up or down. Trendlines were fitted to a first order exponential decay function.

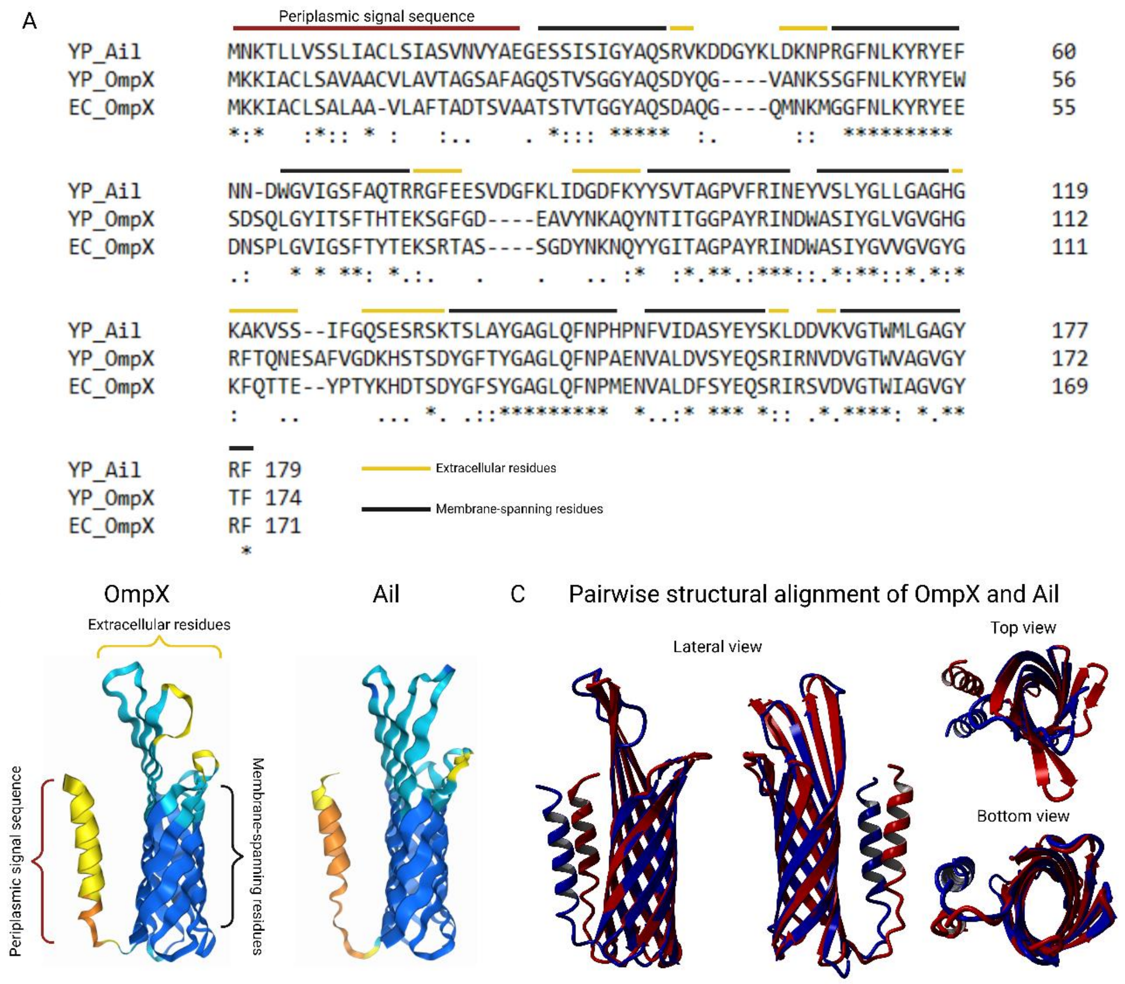

**Supplementary Figure 6. Amino acid and structural comparison of OmpX and Ail in *Y. pseudotuberculosis* YPIII**

(A) The primary amino acid sequence of OmpX (*YPK_1606*) and Ail (*YPK_1268*) as translated from the genome of *Y. pseudotuberculosis* YPIII is shown aligned to *E. coli* OmpX (PDB no. 1QJ8) [3]. Periplasmic signal sequence is denoted by a red bar above the amino acid sequence. Membrane-spanning residues are indicated by a black bar above the sequences and extracellular residues are denoted by a yellow bar. (B) OmpX (ipTM= -; pTM=0.76) and Ail (ipTM= -; pTM=0.76) structures as predicted by AlphaFold3 [4]. Colors denote confidence values for the prediction blue (very high pIDDT > 90), light blue (confident 90> pIDDT >70), yellow (low 70> pIDDT >50), orange (very low pIDDT < 50). (C) Pairwise structural alignment generated by YASARA using the MUltiple STructural AligNment AlGorithm (MUSTANG) [5]. Structural alignment between OmpX (blue) and Ail (red) has a Calpha RMSD of 0.664 Å over 137 aligned residues with 46.72% sequence identity.

**Supporting information – S1 Table**

S1 Table: Bacterial strains and plasmids used in this study.

| Strain or Plasmid | Relevant genotype | Reference |
|---|---|---|
| ***Escherichia coli*** | | |
| **DH5α** | | |
| ***S17-1 λpir*** | RP4-2 Tc::Mu-Km::Tn7 (λpir) | Simon et al., 1983 |
| ***Yersinia pseudotuberculosis*** | | |
| **YPIII** | pIB1, wild type | Bölin et al., 1982 |
| **YPIII ΔCyaR** | pIB1, ΔCyaR::neoR, Kan$^r$ | This study |
| **YPIII Δ*crp*** | pIB1, Δ*crp* | Heroven et al., 2012 |
| **YPIII Δ*csrA::neoR*** | pIB1, Δ*csrA*::*neoR*, Kan$^r$ | Heroven et al., 2008 |
| **YPIII Δ*hfq*** | pIB1, Δ*hfq* | Heroven et al., 2012 |
| **YPIII OmpX::3xFLAG** | pIB1, OmpX::3xFLAG, Cm$^r$ | This study |
| **YPIII ΔCyaR OmpX::3xFLAG** | pIB1, ΔCyaR::*neoR*, OmpX::3xFLAG, Cm$^r$, Kan$^r$ | This study |
| **YPIII Δ*hfq* OmpX::3xFLAG** | pIB1, Δ*hfq*, OmpX::3xFLAG, Cm$^r$ | This study |
| **YPIII ΔOmrA OmpX::3xFLAG** | pIB1, ΔOmrA, OmpX::3xFLAG, Cm$^r$ | Unpublished Data |
| **Plasmids** | | |
| **pBAD2-*bgaB*-His** | reporter gene vector, *bgaB* with His-tag at the C-terminal end, P$_{BAD}$ promoter, *araC*, Ap$^r$ | Klinkert et al., 2012 |
| **pBAD2-*yopN*-*bgaB*-His** | pBAD2-*bgaB*-His, short 5'-UTR of *yopN* (pYP0065) plus 30 bp of the coding region, translational fusion | Pienkoß et al., 2021 |
| **pUC18** | cloning vector, Ap$^r$ | Norrander et al., 1983 |
| **pDM4::3xFLAG** | pDM4, sacBR, oriT, oriR6K, Gm$^r$, Cm$^r$ | This study |
| **pYD32** | pGP704, sacBR, oriT, oriR6K, Ap$^r$, Kan$^r$, integration of neoR inside the CyaR locus for generating YPIII ΔCyaR::neoR | Heroven et al., 2012 |
| **pMA** | colE1, Ap$^r$, Synthetic | Thermo Scientific, Waltham, USA |
| **pXG0** | pXG empty vector, Cm$^r$ | Urban & Vogel 2007 |
| **pXG1** | pXG GFP alone, Cm$^r$ | Urban & Vogel 2007 |
| **pGM930** | pBAD24-Δ1 derivative, pHP45 tΩ terminator downstream of araBp, Ap$^r$ | Delvillani et al., 2014 |
| **pBO5009** | pGM930, 89bp of CyaR (Ysr159), pBAD promoter, araC, Apr, for CyaR complementation | This study |

| | | |
|---|---|---|
| **pBO6885** | pDM4, 1346 bp (782bp upstream of TSS and 564bp downstream of TSS) of *ompX*$^{3x\text{-}FLAG}$ (YPK_1606) without stop codon, Cm$^{r}$ | This study |
| **pBO7535** | pXG, 5'-UTR of *ompX (*YPK_1606) plus 45 bp of the coding region, GFP translational fusion | This study |
| **pBO6899** | pBAD2-*bgaB*-His, 5’-UTR of *ompX* (YPK_1606) plus 30 bp of the coding region, translational fusion | This study |
| **pBO8202** | pBAD2-*bgaB*-His, 5’-UTR of *ompX* (YPK_1606) plus 30 bp of the coding region, R1 variant (UU23-24CC) translational fusion | This study |
| **pBO8203** | pBAD2-*bgaB*-His, 5’-UTR of *ompX* (YPK_1606) plus 30 bp of the coding region, R2 variant (UU23-24CC; G28C) translational fusion | This study |
| **pBO5019** | pUC18, 89bp of CyaR (Ysr159) with T7 promoter sequence, runoff transcription plasmid for structure probing | This study |
| **pBO7428** | pMA, CyaR with T7 promotor sequence, runoff plasmid for structure probing, stable variant (UUU82-83,87GGC), Synthetic | This study |
| **pBO7430** | pMA, CyaR with T7 promotor sequence, runoff plasmid for structure probing, open variant (UUC84-86AAA), Synthetic | This study |
| **pBO7904** | pUC18, 103bp 5'-UTR of *ompX* (YPK_1606) plus 60 bp with T7 promoter sequence, runoff transcription plasmid for structure probing | This study |

**Supporting information – S2 Table**

S2 Table: Oligonucleotides used in this study.

| Primer Name | Sequence 5'-3' | Purpose | Plasmid |
|---|---|---|---|
| **pBAD2-BgaB Constructs** | | | |
| OmpX_BgaB_Fw | TTT*GCTAGC*GTGTTTTTGTGTCACTCAAT | Forward primer to amplify 5'-UTR of *ompX* + 30nt CDS insert translationally fused to BgaB gene for β-Glactosidase assay | pBO6899 |
| OmpX_BgaB_Rv | TTT*GAATTC*TACCGCTGAAAGACATGCA | Reverse primer to amplify 5'-UTR of *ompX* + 30nt CDS translationally fused to BgaB gene for β-Glactosidase assay | |
| OmpX_WT_SD_Fw | TGCGTCTTTCGAGGTGGTTATG | Universal wild type forward for site directed mutagenesis / paired with OmpX_rep1_Rv to create pBO8202 | |
| OmpX_R2_G28C_Fw | TGCCTCTTTCGAGGTGGTTATG | Forward primer to amplify 5'-UTR of *ompX* + 45nt CDS for site directed mutagenesis / paired with OmpX_Rep1_Rv to create pBO8203 | pBO8203 |
| OmpX_Rep1_Rv | GGTTATTGAGTGACACAAAAACAC | Reverse primer to amplify 5'-UTR of *ompX* + 45nt CDS for site directed mutagenesis / paired with Universal wild type to create pBO8202 / paired with OmpX_R2_G28C_Fw to create pBO8203 | pBO8202 |
| **pXG-GFP Constructs** | | | |
| pXG_UTRs_Fw | GCTAGCAAAGGAGAAGAACTTTTCACTG | pair with pXG_Gibson_Rv for vector linearization | |
| pXG_OmpX_gib_Fw | CAGTGATAGAGATACTGAGCACATGTGTTTTTGTGTCACTCAATAATTTGCG | 5'UTR of *ompX* (YPK_1606) up to 15AA | pBO7535 |
| pXG_OmpX_gib_Rv | CAGTGAAAAGTTCTTCTCCTTTGCTAGCTAACACACACGCTGCTACCGC | | |
| pXG_Ev_Gibson_Fw | GAATTCGAGCATTTAAATCTAGAGGC | To create pXG0 true empty vector | pXG0 |
| pXG_Ev_Gibson_Rv | ATGTGCTCAGTATCTCTATCACTG | | |
| pXG1_GFPEv_Fw | GAGGAGAAAGGTACCATGGCTAGCAAAGGAGAAGAACTTTTCACTGGAG | To recreate pXG1 Empty vector | pXG1 |
| pXG1_GFPEv_Rv | TTTAATGAATTCGGTCAGTGCGTCCTGCTGATGTGCTCAGTATCTCTATCACTG | | |
| **pUC18** | | | |
| OmpX_rnf_Fw | A*GAAATTAATACGACTCACTATAGGG*GTGTTTTTGTGTCACTCAATAATTTGCG | 5'-UTR of *ompX* + 60nt CDS for structure probing | pBO7904 |
| OmpX_rnf_Rv | AAGATATCACCTGCGGTCACGGCTAACA | 5'-UTR of *ompX* + 60nt CDS for structure probing / Reverse primer used for primer extension inhibition assays | |
| CyaR_T7_Fw | *GAAATTAATACGACTCACTATAGGG*GAGTACAATCAATAACTAAAAAAGTG | *Y.pseudotuberculosis* cyaR fw with T7 promoter for cloning into pUC18, runoff | pBO5019 |
| CyaR_EcoRV_Rv | TTGATATCAAAGAAAAGGCCAGCCCAAG | *Y.pseudotuberculosis* cyaR rv for cloning into pUC18, runoff | |
| pGM930 | | | |
| CyaR_PstI_Rv | TTCTGCAGAAAGAAAAGGCCAGCCCAAG | *Y.pseudotuberculosis* cyaR rv for cloning into pGM930 | pBO5009 |
| CyaR_NcoI_Fw | TTCCATGGGAGTACAATCAATAACTAAAAAAGTG | *Y.pseudotuberculosis* cyaR fw for riboprobe and cloning into pGM930-NcoI | |
| **Chromosomal integration strains** | | | |
| ompX_SacI_Fw | TTTGAGCTCCCAGTGGTAACAGGAACCAT | To construct the ompX::3xFLAG for chromosomal integration | pBO6885 |
| ompX_XbaI_Rv | TTTTCTAGAGAAAGTGTAACCTACGCCAGC | To construct the ompX::3xFLAG for chromosomal integration | |
| CyaR_internal_Fw | TTAAGCTTTGCCACAGATAAAGTGGCATT | To validate ΔCyaR strain | |
| CyaR_internal_Rv | TTGGATCCCTGTTGGGATAATTATTATTTTAGG | To validate ΔCyaR strain | |
| CyaR_external_Fw | TTTGACCGCTTCGCGGAC | To validate ΔCyaR strain | |
| CyaR_external_Rv | ATGTTGAACCGACGCTGGG | To validate ΔCyaR strain | |
| **RNA Probes** | | To validate ΔCyaR strain | |
| CyaR_probe_Fw | TTCCATGGGAGTACAATCAATAACTAAAAAAGTG | forward primer for the synthesis of a CyaR probe | |
| CyaR_probe_Rv | *GAAATTAATACGACTCACTATAGGG*AAAGAAAAGGCCAGCCCAAGGG | reverse primer for the synthesis of a CyaR probe | |
| ompX_probe_Fw | TTTATGCATGTGTTTTTGTGTCACTCAATAATT | forward primer for the synthesis of an *ompX* probe | |
| ompX_probe_Rv | *GAAATTAATACGACTCACTATAGGG*ACCAGAAACGGTGCTTTGAC | reverse primer for the synthesis of an *ompX* probe | |
| 5S_probe_Fw | CTGGCGGCCATAGCGCGG | forward primer for the synthesis of a 5S rRNA probe | |
| 5S_probe_Rv | TTT*GAAATTAATACGACTCACTATAGGG*CCTGGCAGTGTCCTACTCTCG | reverse primer for the synthesis of a 5S rRNA probe | |
